# Insulin-stimulated glucose uptake partly relies on p21-activated kinase (PAK)-2, but not PAK1, in mouse skeletal muscle

**DOI:** 10.1101/543736

**Authors:** Lisbeth L. V. Møller, Merna Jaurji, Rasmus Kjøbsted, Giselle A. Joseph, Agnete B. Madsen, Jonas R. Knudsen, Annemarie Lundsgaard, Nicoline R. Andersen, Peter Schjerling, Thomas E. Jensen, Robert S. Krauss, Erik A. Richter, Lykke Sylow

## Abstract

**Objective:** Skeletal muscle glucose uptake is essential for maintaining whole-body glucose homeostasis and accounts for the majority of glucose disposal in response to insulin. The group I p21-activated kinase (PAK) isoforms PAK1 and PAK2 are activated in response to insulin in skeletal muscle. Interestingly, PAK1/2 signalling is impaired in insulin-resistant mouse and human skeletal muscle and PAK1 has been suggested to be required for insulin-stimulated GLUT4 translocation. However, the relative contribution of PAK1 and PAK2 to insulin-stimulated glucose uptake in mature skeletal muscle is unresolved. The aim of the present investigation was to determine the requirement for PAK1 and PAK2 in whole-body glucose homeostasis and insulin-stimulated glucose uptake in skeletal muscle.

**Methods:** Glucose uptake was measured in isolated skeletal muscle incubated with a pharmacological inhibitor (IPA-3) of group I PAKs and in muscle from whole-body PAK1 knockout (KO), muscle-specific PAK2 (m)KO and double whole-body PAK1 and muscle-specific PAK2 knockout mice.

**Results:** The whole-body respiratory exchange ratio was largely unaffected by lack of PAK1 and/or PAK2. Whole-body glucose tolerance was mildly impaired in PAK2 mKO, but not PAK1 KO mice. IPA-3 partially reduced (−20%) insulin-stimulated glucose uptake in mouse soleus muscle. In contrast to a previous study of GLUT4 translocation in PAK1 KO mice, PAK1 KO muscles displayed normal insulin-stimulated glucose uptake *in vivo* and in isolated muscle. On the contrary, glucose uptake was slightly reduced in response to insulin in glycolytic extensor digitorum longus muscle lacking PAK2, alone (−18%) or in combination with PAK1 KO (−12%).

**Conclusions:** Insulin-stimulated glucose uptake partly relies on PAK2, but not PAK1, in mouse skeletal muscle. Thus, the present study challenges that group I PAKs, and especially PAK1, are major regulators of whole-body glucose homeostasis and insulin-stimulated glucose uptake in skeletal muscle.

## 1. Introduction

Skeletal muscles account for the majority of insulin-mediated whole-body glucose disposal [1, 2] and muscle insulin resistance is an early defect in the pathophysiology of peripheral insulin resistance and type 2 diabetes mellitus [1, 3]. As diabetes globally is approaching epidemic proportions, it is important to understand the mechanisms regulating glucose uptake by skeletal muscle.

Insulin stimulates glucose uptake in skeletal muscle by activation of a signalling cascade that leads to the translocation of glucose transporter (GLUT)4-containing vesicles to the sarcolemma and transverse tubuli [4]. This signalling cascade has been proposed to include activation of p21-activated kinase 1 (PAK1) downstream of PI3K [5–7]. PAKs are serine/threonine kinases and involved in numerous signalling networks regulating essential cellular activities, including cell proliferation, differentiation, apoptosis, and cytoskeleton dynamics [8–11]. Group I PAKs (PAK1-3) are downstream targets of the Rho GTPases Cdc42 and Rac1 [12]. Previous studies suggest that only PAK1 and PAK2 are expressed in skeletal muscle, whereas PAK3 mRNA and protein expression is below the detection limit [6,13,14]. In muscle cells and mouse skeletal muscle, PAK1 is proposed to be required for GLUT4 translocation in response to insulin stimulation [6, 7], likely downstream of Rac1 [15–20]. Thus, together with Akt, proposed to regulate GLUT4 translocation via phosphorylation of the Rab GTPase-activating protein TBC1D4 [21–23], Rac1-PAK1 activation is suggested to be necessary for insulin-stimulated GLUT4 translocation.

Upon activation of PAK1 and PAK2, conformational changes allow autophosphorylation of T423 and T402, respectively, thereby relieving the autoinhibition of PAK1 and PAK2 [12,24,25]. In vastus lateralis muscle from subjects with obesity and type 2 diabetes, phosphorylation of PAK1/2 at T423/402 in response to insulin was 50% reduced compared to healthy controls [18]. Likewise, insulin-stimulated pPAK1/2 T423/402 was diminished in palmitate-treated insulin-resistant L6 myotubes, even though upstream of PAK1/2, insulin-stimulated Rac1-GTP binding (i.e. activation) was not impaired [26]. Together, these studies [18, 26] associate dysregulated activity of PAK1 and PAK2 with insulin resistance. In addition, a pharmacological inhibitor of group I PAKs, IPA-3 abolished insulin-stimulated GLUT4 translocation and glucose uptake in L6 myoblasts and myotubes overexpressing myc-tagged GLUT4 (L6-GLUT4myc), respectively [6]. This indicates that group I PAKs are required for insulin-stimulated glucose uptake. The effect of IPA-3 has largely been ascribed to inhibition of PAK1, as whole-body genetic ablation of PAK1 in mice impaired glucose tolerance [7, 27] and blocked insulin-stimulated GLUT4 translocation in skeletal muscle [7]. Further supporting PAK1 being the major PAK isoform regulating GLUT4 translocation, insulin-stimulated GLUT4 translocation was unaffected by a 75% knockdown of PAK2 in L6-GLUT4myc myoblasts [6]. The suggested downstream mechanisms whereby PAK1 regulates GLUT4 translocation include simultaneous cofilin-mediated actin depolymerization and N-WASP-cortactin-mediated actin polymerization [6,28,29].

Although such studies implicate group I PAKs, and in particular PAK1, in the regulation of glucose homeostasis and GLUT4 translocation in skeletal muscle, the relative role of PAK1 and PAK2 in insulin-stimulated glucose uptake remains to be identified in mature skeletal muscle. Therefore, we performed a systematic series of pharmacologic and genetic experiments to analyze the involvement of group I PAKs in the regulation of insulin-stimulated glucose uptake in mouse skeletal muscle. We hypothesized that group I PAKs, and in particular PAK1, would be necessary for glucose uptake in response to insulin. Contradicting our hypothesis, our results revealed that insulin-stimulated glucose uptake partly relies on PAK2 in glycolytic mouse muscle, while PAK1 is dispensable for whole-body glucose homeostasis and insulin-stimulated muscle glucose uptake.

## 2. Methods

### 2.1. Animals

All animal experiments complied with the European Convention for the protection of vertebrate animals used for experimental and other scientific purposes (No. 123, Strasbourg, France, 1985; EU Directive 2010/63/EU for animal experiments) and were approved by the Danish Animal Experimental Inspectorate. All mice were maintained on a 12:12-hour light-dark cycle and housed at 22°C (with allowed fluctuation of ±2°C) with nesting material. The mice were group-housed. Female C57BL/6J mice (Taconic, Denmark) were used for the inhibitor incubation study. The mice received a standard rodent chow diet (Altromin no. 1324; Brogaarden, Denmark) and water ad libitum.

#### 2.1.1. Whole-body PAK1^−/−^ mice

For a complete overview of the different cohorts of genetically modified mice, see supplementary Table S1. Whole-body PAK1^−/−^ mice on a C57BL/6J background were generated as previously described [30]. The mice were obtained by heterozygous crossing. PAK1^−/−^ mice (referred to as PAK1 KO) and paired littermate PAK1^+/+^ mice (referred to as controls) were used for experiments. Female and male mice were used for measurements of body composition, glucose tolerance and insulin-stimulated glucose uptake in isolated muscle. The mice were 12-24 weeks of age at the time of tissue dissection and measurement of glucose uptake. Number of mice in each group: Control, *n = 6/7* (female/male); PAK1 KO, *n = 4/8*. Mice received standard rodent chow diet and water ad libitum. For measurement of *in vivo* insulin-stimulated glucose uptake in chow- and 60E% high-fat diet (HFD; no. D12492; Brogaarden, Denmark)-fed PAK1 KO mice, mice were assigned to a chow or HFD group. Chow-fed mice were 10-24 weeks of age at the time of glucose uptake measurements and tissue dissection. Number of mice in each group: Control-Chow, *n = 14/8* (female/male); PAK1 KO-Chow, *n = 6/4*. For mice receiving HFD, the diet intervention started at 6-16 weeks of age and lasted for 21 weeks. HFD-fed mice were used for body composition, glucose tolerance and *in vivo* glucose uptake and were 27-37 weeks of age at the time of glucose uptake measurements and tissue dissection. Number of mice in each group: Control-HFD, *n = 7/7* (female/male); PAK1 KO-HFD, *n = 11/5*. Energy intake was measured over a period of 10 weeks in another cohort of mice. Number of mice in each group: Chow, *n = 8/8* (Control/PAK1 KO); HFD, *n = 8/8*. Mice had access to their respective diet and water ad libitum.

#### 2.1.2. Double PAK1^−/−^;PAK2^fl/fl^;MyoD^iCre/+^ mice

Double knockout mice with whole-body knockout of PAK1 and conditional, muscle-specific knockout of PAK2, PAK1^−/−^;PAK2^fl/fl^;MyoD^iCre/+^ were generated as previously described [13]. The mice were on a mixed C57BL/6/FVB background. PAK1^−/−^;PAK2^fl/fl^;MyoD^iCre/+^ were crossed with PAK1^+/-^;PAK2^fl/fl^;MyoD^+/+^ to generate littermate PAK1^−/−^;PAK2^fl/fl^;MyoD^iCre/+^ (referred to as 1/m2 dKO), PAK1^−/−^;PAK2^fl/fl^;MyoD^+/+^ (referred to as PAK1 KO), PAK1^+/-^;PAK2^fl/fl^;MyoD^iCre/+^ (referred to as PAK2 mKO), and PAK1^+/-^;PAK2^fl/fl^;MyoD^+/+^ (referred to as controls) used for experiments. Female and male mice were used for measurement of insulin-stimulated glucose uptake in isolated muscle. The mice were 10-16 weeks of age at the time of tissue dissection and glucose uptake measurements. Number of mice in each group: Control, *n = 6/4* (female/male); PAK1 KO, *n = 5/4*, PAK2 mKO, *n = 6/4*, 1/m2 dKO, *n = 6/3*. Another cohort of mice was used for whole-body metabolic measurements. The first measurement (insulin tolerance) was at 11-24 weeks of age and the last measurement was at 23-33 weeks of age (home cage calorimetry). Number of mice in each group: Control, *n = 9/11* (female/male); PAK1 KO, *n = 8/10*, PAK2 mKO, *n = 12/9*, 1/m2 dKO, *n = 9/14*. For some of the metabolic measurements, only a subgroup of mice was used as indicated in the relevant figure legends. Mice received standard rodent chow diet and water ad libitum.

### 2.2. Body composition

Body composition was analyzed using magnetic resonance imaging (EchoMRI-4in1TM, Echo Medical System LLC, Texas, USA). Chow-fed PAK1 KO and control littermates were assessed at 7-19 weeks of age. HFD-fed PAK1 KO and control littermates were assessed 18-19 weeks into the diet intervention (24-34 weeks of age). Chow-fed PAK1 KO, PAK2 mKO, 1/m2 dKO mice and control littermates were assessed at 16-29 weeks of age.

### 2.3. Glucose tolerance test (GTT)

Glucose tolerance was assessed in 8-20 weeks of age chow-fed PAK1 KO mice and in week 14 of the diet intervention of HFD-fed PAK1 KO mice (20-30 weeks of age). In chow-fed PAK1 KO, PAK2 mKO, 1/m2 dKO mice and control littermates, glucose tolerance was assessed at 13-26 weeks of age. Prior to the test, chow- and HFD-fed PAK1 KO mice and control littermates fasted for 12 hours from 10 p.m, while chow-fed PAK1 KO, PAK2 mKO, 1/m2 dKO mice and control littermates fasted for 6 hours from 6 a.m. D-mono-glucose (2 g kg^−1^ body weight) was administered intraperitoneal (i.p) and blood was collected from the tail vein and blood glucose concentration determined at the indicated time points using a glucometer (Bayer Contour, Bayer, Switzerland). Incremental Area Under the Curve (AUC) from the basal blood glucose concentration was determined using the trapezoid rule. For measurement of plasma insulin, glucose was administered i.p. on a separate experimental day (1-2 weeks after the GTT) and blood was sampled at time points 0 and 20 minutes, centrifuged and plasma was quickly frozen in liquid nitrogen and stored at −20°C until processing. Plasma insulin was analyzed in duplicate (Mouse Ultrasensitive Insulin ELISA, #80-INSTRU-E10, ALPCO Diagnostics, USA). Homeostatic model assessment of insulin resistance (HOMA-IR) was calculated according to the formula: Fasting plasma insulin (mU L^−1^) X Fasting blood glucose (mM)/22.5.

### 2.4. Insulin tolerance test (ITT)

Insulin tolerance was assessed in 11-24 weeks of age chow-fed PAK1 KO, PAK2 mKO, 1/m2 dKO mice and control littermates. Prior to the test, chow-fed PAK1 KO, PAK2 mKO, 1/m2 dKO mice and control littermates fasted for 4 hours from 6 a.m. Insulin (0.5 U kg^−1^ body weight) was administered i.p. and blood was collected from the tail vein and blood glucose concentration determined using a glucometer (Bayer Contour, Bayer, Switzerland) at time point 0, 15, 30, 60, 90 and 120 minutes. For two female control mice and four female PAK2 mKO mice, the ITT had to be stopped before the 120’-time point due to hypoglycemia (blood glucose <1.2 mM). Thus, blood glucose was not measured in these mice for the last couple of time points.

### 2.5. Home cage indirect calorimetry

One week prior to the calorimetric measurements, chow-fed PAK1 KO, PAK2 mKO, 1/m2 dKO mice and control littermates were single-housed in specialized cages for indirect gas calorimetry but uncoupled from the PhenoMaster indirect calorimetry system (TSE PhenoMaster metabolic cage systems, TSE Systems, Germany). After a 2-day acclimation period coupled to the PhenoMaster indirect calorimetry system, oxygen consumption, CO_2_ production, habitual activity (beam breaks) and food intake were measured for 72 hours (TSE LabMaster V5.5.3, TSE Systems, Germany). On day 2, mice fasted during the dark period followed by refeeding on day 3. Respiratory exchange ratio (RER) was calculated as the ratio between CO_2_ production and oxygen consumption. The mice were 23-33 weeks of age.

### 2.6. Incubation of isolated muscles

Soleus and extensor digitorum longus (EDL) muscles were dissected from anaesthetized mice (6 mg pentobarbital sodium 100 g^−1^ body weight i.p.) and suspended at resting tension (4-5 mN) in incubations chambers (Multi Myograph System, Danish Myo Technology, Denmark) in Krebs-Ringer-Henseleit buffer with 2 mM pyruvate and 8 mM mannitol at 30°C, as described previously [31]. Additionally, the Krebs-Ringer-Henseleit buffer was supplemented with 0.1% BSA (v/v). Isolated muscles from female C57BL/6J mice were pre-incubated with 40 µM IPA-3 (Sigma-Aldrich) or as a control DMSO (0.25%) for 25 minutes followed by 30 minutes of insulin stimulation (60 nM; Actrapid, Novo Nordisk, Denmark). Isolated muscles from chow-PAK1 KO were pre-incubated for 30 minutes followed by 30 minutes of insulin stimulation (0.6 nm or 60 nM). Isolated muscles from chow-fed PAK1 KO, PAK2 mKO, 1/m2 dKO mice or control littermates were pre-incubated for 20 minutes followed by 20 minutes of insulin stimulation (60 nM). 2-Deoxyglucose (2DG) uptake was measured together with 1 mM 2DG during the last 10 min of the insulin stimulation period using 0.60-0.75 µCi mL^−1^ [^3^H]-2DG and 0.180-0.225 µCi mL^−1^ [^14^C]-mannitol radioactive tracers as described previously [31]. Tissue-specific [^3^H]-2DG accumulation with [^14^C]-mannitol as an extracellular marker was determined as previously described [32].

### 2.7. In vivo insulin-stimulated 2-Deoxyglucose uptake in PAK1 KO mice during a r.o. ITT

To determine 2DG uptake in skeletal muscle of PAK1 KO mice and littermate controls, [^3^H]-2DG (Perkin Elmer) was administered retro-orbitally (r.o.) in a bolus of saline containing 66.7 μCi mL^−1^ [^3^H]-2DG (∼32.4 Ci/mmol) corresponding to ∼10 μCi/mouse in chow-fed mice or ∼15 μCi/mouse in HFD-fed mice (6 μL g^−1^ body weight) as described [20]. The [^3^H]-2DG saline bolus was with or without insulin (Actrapid, Novo Nordisk, Denmark). Decreased insulin clearance has been observed by us [20] and others in obese rodent [33, 34] and human [35] models. Therefore, to correct for changes in insulin clearance, 0.5 U kg^−1^ body weight of insulin was administered in chow-fed mice whereas only 60% of this dosage was administered to HFD-fed mice. Prior to stimulation, mice fasted for 4 hours from 07:00 and were anaesthetized (7.5/9 mg [Chow/HFD] pentobarbital sodium 100 g^−1^ body weight i.p.) 15 minutes before the r.o. injection. Blood samples were collected from the tail vein after the r.o. injection and analyzed for glucose concentration using a glucometer (Bayer Contour, Bayer, Switzerland) at the time points 0, 5 and 10 minutes. After 10 minutes, skeletal muscle (gastrocnemius, quadriceps, triceps brachii and soleus) were excised, rinsed in saline, and quickly frozen in liquid nitrogen and stored at −80°C until processing. Blood was collected by punctuation of the heart, centrifuged and plasma was quickly frozen in liquid nitrogen and stored at −80°C until processing. Plasma samples were analyzed for insulin concentration and specific [^3^H]-2DG activity. Plasma insulin was analyzed in duplicate (Mouse Ultrasensitive Insulin ELISA, #80-INSTRU-E10, ALPCO Diagnostics, USA). Tissue-specific 2DG-6-phosphate accumulation was measured as described [36, 37]. To determine 2DG clearance from the plasma into the specific tissue, tissue-specific [^3^H]-2DG-6-P was divided by AUC of the plasma-specific [^3^H]-2DG activity at the time points 0 and 10 minutes. To estimate tissue-specific glucose uptake (glucose uptake index), clearance was multiplied by the average blood glucose levels for the time points 0, 5, and 10 minutes. Tissue-specific 2DG clearance and glucose uptake were related to analyzed muscle tissue weight and time.

### 2.8. Muscle molecular analysis

Prior to homogenization, gastrocnemius, quadriceps, and triceps brachii muscles were pulverized in liquid nitrogen. All muscle were homogenized 2 x 30 sec at 30 Hz using a Tissuelyser II (Qiagen, USA) in ice-cold homogenization buffer (10% (v/v) Glycerol, 1% (v/v) NP-40, 20 mM Na-pyrophosphate, 150 mM NaCl, 50 mM HEPES (pH 7.5), 20 mM β-glycerophosphate, 10 mM NaF, 2mM PMSF, 1 mM EDTA (pH 8.0), 1 mM EGTA (pH 8.0), 2 mM Na3VO4, 10 µg mL^−1^ Leupeptin, 10 µg mL^−1^ Aprotinin, 3 mM Benzamidine). After rotation end-over-end for 30 min at 4°C, lysate supernatants were collected by centrifugation (10,854-15,630 x g) for 15-20 min at 4°C.

#### 2.8.1. Immunoblotting

Lysate protein concentration was determined using the bicinchoninic acid method using bovine serum albumin (BSA) standards and bicinchoninic acid assay reagents (Pierce). Immunoblotting samples were prepared in 6x sample buffer (340 mM Tris (pH 6.8), 225 mM DTT, 11% (w/v) SDS, 20% (v/v) Glycerol, 0.05% (w/v) Bromphenol blue). Protein phosphorylation (p) and total protein expression were determined by standard immunoblotting technique loading equal amounts of protein. The polyvinylidene difluoride membrane (Immobilon Transfer Membrane; Millipore) was blocked in Tris-Buffered Saline with added Tween20 (TBST) and 2% (w/v) skim milk or 5% (w/v) BSA protein for 15 minutes at room temperature, followed by incubation overnight at 4°C with a primary antibody (Table 1). Next, the membrane was incubated with horseradish peroxidase-conjugated secondary antibody (Jackson Immuno Research) for 1 hour at room temperature. Bands were visualized using Bio-Rad ChemiDocTM MP Imaging System and enhanced chemiluminescence (ECL+; Amersham Biosciences). Densitometric analysis was performed using Image LabTM Software, version 4.0 (Bio-Rad, USA; RRID:SCR_014210). Coomassie brilliant blue staining was used as a loading control [38].

**Table 1.** Antibody Table

| **Antibody name** | **Antibody ID (RRID)** | **Manufacturer; Catalog Number;** | **Species Raised in; Monoclonal or Polyclonal** | **Antibody dilution** | **Blocking buffer** |
| --- | --- | --- | --- | --- | --- |
| Akt2 | AB_2225186 | Cell Signaling Technology; 3063 | Rabbit; Monoclonal antibody | 1:1000 | 2% milk |
| pAkt S473 | AB_329825 | Cell Signaling Technology; 9271 | Rabbit; Polyclonal antibody | 1:1000 | 2% milk |
| pAkt T308 | AB_329828 | Cell Signaling Technology; 9275 | Rabbit; Polyclonal antibody | 1:1000 | 2% milk |
| GLUT4 | AB_2191454 | Thermo Fisher Scientific; PA1-1065 | Rabbit; Polyclonal antibody | 1:1000 | 2% milk |
| PAK1 | AB_330222 | Cell Signaling Technology; 2602 | Rabbit; Polyclonal antibody | 1:1000 | 2% milk |
| PAK2 | AB_2283388 | Cell Signaling Technology; 2608 | Rabbit; Polyclonal antibody | 1:1000 | 2% milk |
| pPAK1/2 T423/402 | AB_330220 | Cell Signaling Technology; 2601 | Rabbit; Polyclonal antibody | 1:1000 | 5% BSA |
| TBC1D4 | AB_492639 | Millipore; 07-741 | Rabbit; Polyclonal antibody | 1:1000 | 2% milk |
| pTBC1D4 T642 | AB_2651042 | Cell Signaling Technology; 8881 | Rabbit; Monoclonal antibody | 1:1000 | 2% milk |

### 2.9. Statistical analyses

Data are presented as mean ± S.E.M. or when applicable mean ± S.E.M. with individual data points shown. Statistical tests varied according to the dataset being analyzed and the specific tests used are indicated in the figure legends. Datasets were normalized by square root, log10, inverse or inverse square root transformation if not normally distributed or unequal equal variance. If the null hypothesis was rejected, Tukey’s post hoc test was used to evaluate significant main effects of genotype and significant interactions in ANOVAs. P < 0.05 was considered statistically significant. P<0.1 was considered a tendency. Except for mixed-effects model analyses performed in GraphPad Prism, version 8.2.1. (GraphPad Software, La Jolla, CA, USA; RRID:SCR_002798), all statistical analyses were performed using Sigma Plot, version 13 (Systat Software Inc., Chicago, IL, USA; RRID:SCR_003210). Due to missing values ascribed to hypoglycemia, differences between genotypes and the effect of insulin administration were assessed with a mixed-effects model analysis in Fig. 5F and S5F.

## 3. Results

### 3.1. Pharmacological inhibition of group I PAKs partially reduces insulin-stimulated glucose uptake in mouse soleus muscle

To investigate the involvement of group I PAKs in insulin-stimulated glucose uptake in mouse skeletal muscle, we first analyzed 2DG uptake in isolated soleus and EDL muscle in the presence of a pharmacological inhibitor, IPA-3. While glucose uptake *in vivo* is influenced by glucose delivery, GLUT4 translocation and muscle metabolism [39], glucose delivery is constant in isolated skeletal muscle and surface membrane GLUT4 is the limiting factor [40–42]. Therefore, 2DG uptake in isolated muscles likely reflects GLUT4 translocation. IPA-3 is a well-characterized inhibitor of group I PAKs (PAK1-3) [6, 43] and shown to completely abolish insulin-stimulated GLUT4 translocation and glucose uptake in L6-GLUT4myc myoblasts and myotubes, respectively [6]. In soleus, 2DG uptake increased 4.5-fold upon maximal insulin-stimulation, an increase that was partly reduced (−20%) in IPA-3-treated muscles (Fig. 1A). IPA-3 did not significantly (p=0.080) impair insulin-stimulated (+2.4-fold) 2DG uptake in EDL muscle (Fig. 1B). We confirmed that IPA-3 treatment abolished insulin-stimulated phosphorylation of (p)PAK1 T423 in both soleus and EDL muscle (Fig. 1C+D). In contrast, insulin-stimulated pAkt T308 (Fig. 1E+F) and pAkt S473 (Fig. 1G+H) were unaffected by IPA-3 treatment, suggesting that IPA-3 did not interfere with insulin signalling to Akt. Altogether this suggests that group I PAKs are only partially involved in insulin-stimulated glucose uptake in isolated mouse muscle.

**Figure 1:**
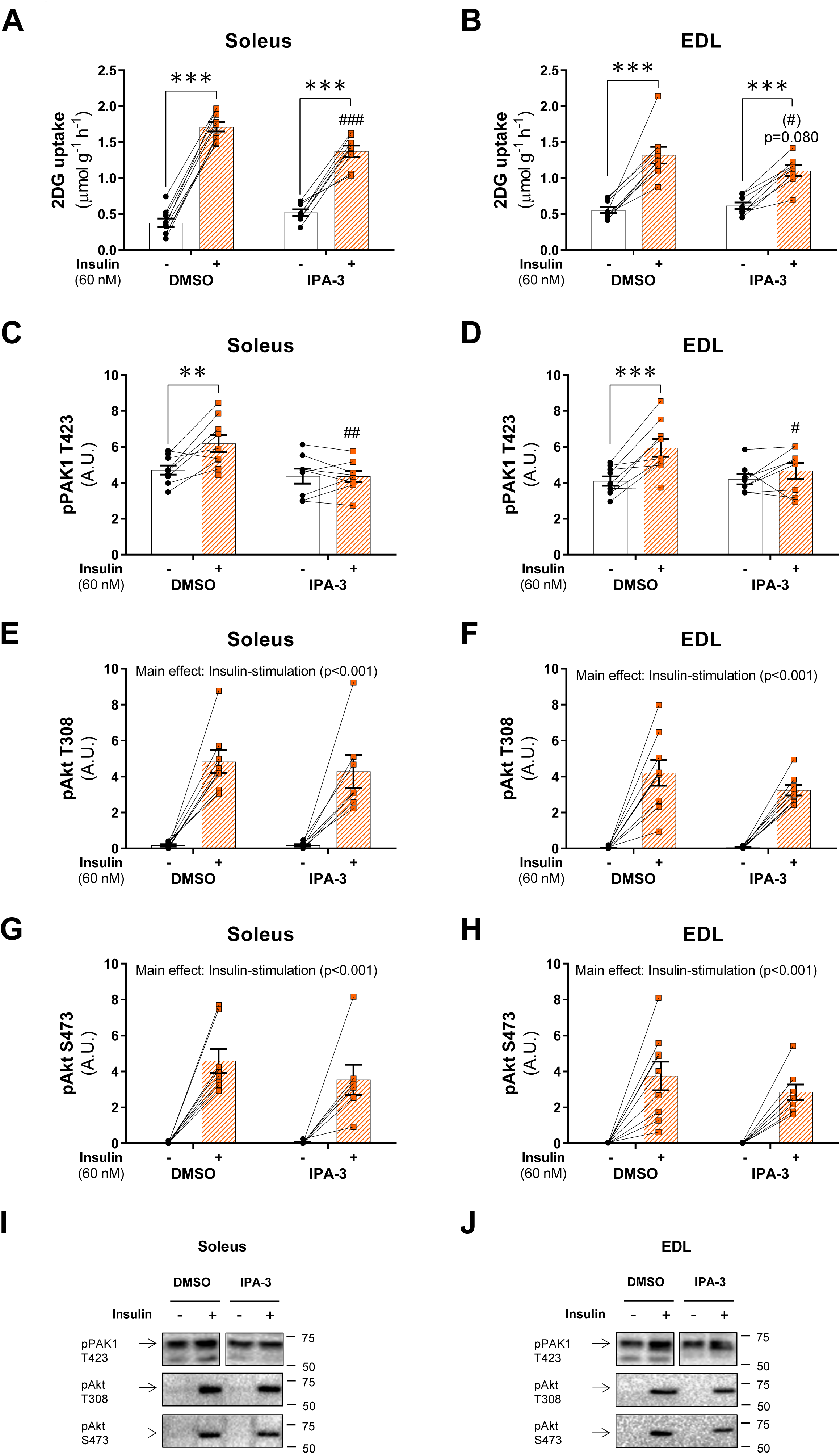
Pharmacological inhibition of group I PAKs partially reduces insulin-stimulated glucose uptake in mouse soleus muscle. **(A-B)** Insulin-stimulated (60 nM) 2-Deoxyglucose (2DG) uptake in isolated soleus (A) and extensor digitorum longus (EDL, B) muscle ± 40 µM IPA-3 or as a control DMSO (0.25%). Isolated muscles were pre-incubated for 25 minutes followed by 30 minutes of insulin stimulation with 2DG uptake analyzed for the final 10 minutes of stimulation. **(C-H)** Quantification of phosphorylated (p)PAK1 T423, pAkt T308, and pAkt S473 in insulin-stimulated soleus (C, E, and G) and EDL (D, F, and H) muscle ± 40 µM IPA-3 or as a control DMSO (0.25%). Some of the data points were excluded due to the quality of the immunoblot, and the number of determinations was *n = 8/7* for pAkt S473 and T308 in soleus muscle. **(I-J)** Representative blots showing pPAK1 T423, pAkt T308, and pAkt S473 in soleus (I) and EDL (J) muscle. Statistics were evaluated with a two-way repeated measures (RM) ANOVA. Main effects are indicated in the panels. Interactions in two-way RM ANOVA were evaluated by Tukey’s post hoc test: Insulin stimulation vs. basal **/*** (p<0.01/0.001); IPA-3 vs. DMSO (#)/#/##/### (p<0.1/0.05/0.01/0.001). Unless otherwise stated previously in the figure legend, the number of determinations in each group: Soleus, *n = 9/8* (DMSO/IPA-3); EDL, *n = 9/8*. Data are presented as mean ± S.E.M. with individual data points shown. Paired data points are connected with a straight line. A.U., arbitrary units.

### 3.2. PAK1 knockout does not affect whole-body glucose tolerance or insulin-stimulated glucose uptake in isolated skeletal muscle

We next sought to confirm our findings in mice with whole-body lack of the PAK1 isoform (PAK1 KO) and therefore a complete knockout of PAK1 in skeletal muscle (Fig. 2A). In chow-fed mice, fat mass, lean body mass, body weight and energy intake (Fig. 2B-C) were similar between whole-body PAK1 KO and littermate controls, as also reported previously [44]. During a GTT, the lack of PAK1 had no effect on the blood glucose response (Fig. 2D-E) or plasma insulin concentration (Fig. 2F). Additionally, HOMA-IR, a measure of basal glucose homeostasis (Fig. 2G), and both submaximal and maximal insulin-stimulated 2DG uptake in isolated soleus and EDL muscle were unaffected by PAK1 KO (Fig. 2H-I). Thus, unexpectedly, genetic ablation of PAK1 alone did not impair whole-body glucose tolerance, or skeletal muscle insulin sensitivity (submaximal insulin-stimulated glucose uptake) or insulin responsiveness (maximal insulin-stimulated glucose uptake) in divergence to previous reports [6,7,11].

**Figure 2:**
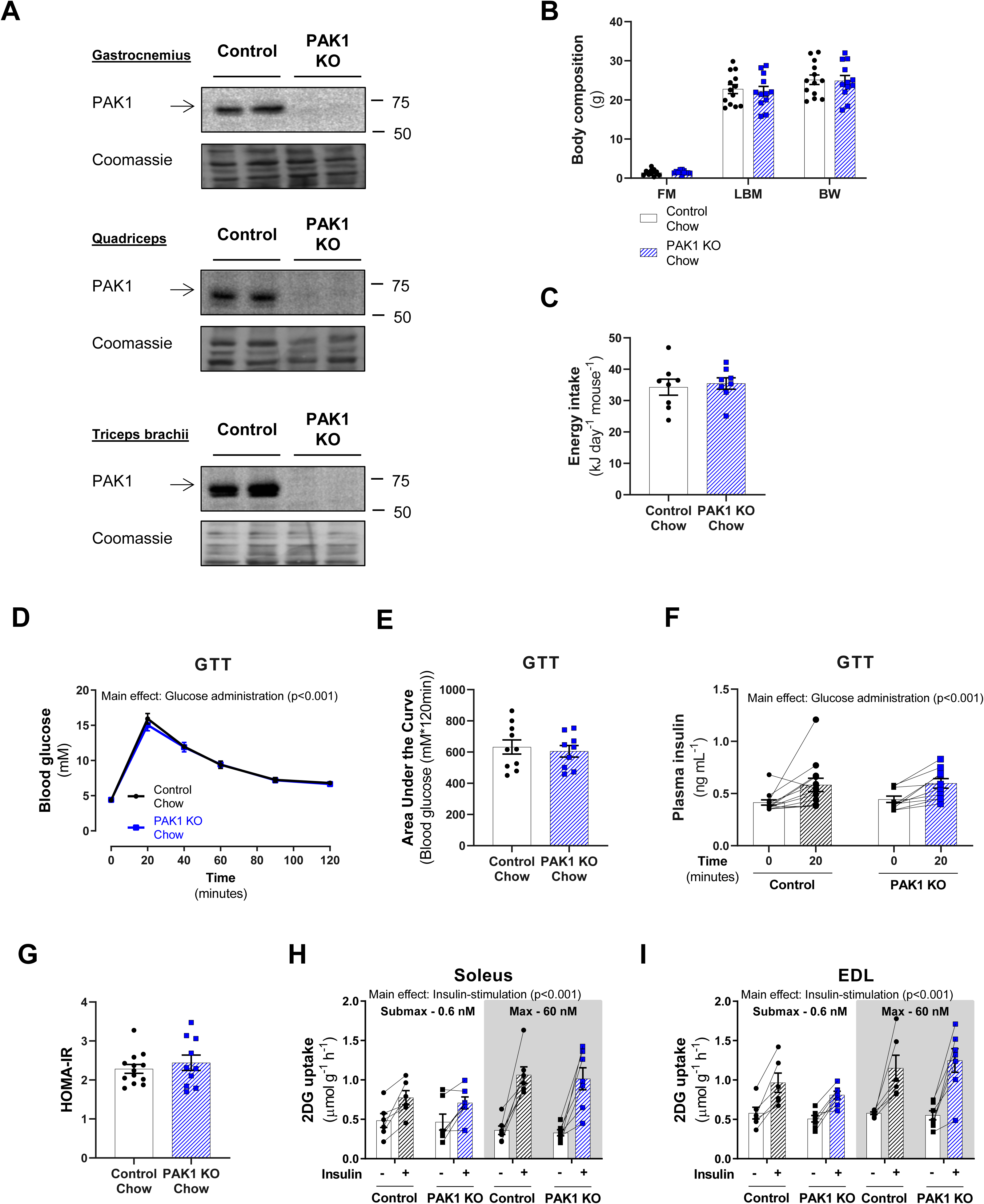
PAK1 knockout does not affect whole-body glucose tolerance or insulin-stimulated glucose uptake in isolated skeletal muscle. **(A)** Representative blots showing PAK1 protein expression in gastrocnemius, quadriceps, and triceps brachii muscle from PAK1 knockout (KO) mice or control littermates. **(B)** Body composition (FM: Fat mass; LBM: Lean body mass; BW: Body weight) in gram in chow-fed PAK1 KO mice (*n = 12*) or control littermates (*n = 13*). The mice were 7-19 weeks of age. Statistics were evaluated with a Student’s t-test. **(C)** Energy intake in chow-fed PAK1 KO mice (*n = 8*) or control littermates (*n = 8*). Energy intake was monitored in a separate cohort of mice. Statistics were evaluated with a Student’s t-test. **(D)** Blood glucose levels during a glucose tolerance test (GTT) in chow-fed PAK1 KO mice (*n = 9*) or control littermates (*n = 10*). The mice were 8-20 weeks of age. Statistics were evaluated with a two-way repeated measures (RM) ANOVA. **(E)** Incremental Area Under the Curve (AUC) for blood glucose levels during the GTT in panel D. Statistics were evaluated with a Student’s t-test. **(F)** Plasma insulin values during a GTT in chow-fed PAK1 KO mice (*n = 10*) or control littermates (*n = 13*). The mice were 9-21 weeks of age. Statistics were evaluated with a two-way RM measures ANOVA. **(G)** Homeostatic Model Assessment of Insulin Resistance (HOMA-IR) in chow-fed PAK1 KO mice (*n=10*) or control littermates (*n=13*). Statistics were evaluated with a Student’s t-test. **(H-I)** Submaximal (0.6 nM) and maximal (60 nM) insulin-stimulated 2-Deoxyglucose (2DG) uptake in isolated soleus (H) and extensor digitorum longus (EDL; I) muscle from PAK1 KO mice or littermate controls. Isolated muscles were pre-incubated for 30 minutes followed by 30 minutes of insulin stimulation with 2DG uptake analyzed for the final 10 minutes of stimulation. The mice were 12-24 weeks of age. The number of determinations in each group: Soleus-Control, *n = 6/7* (Submax/max); Soleus-PAK1 KO, *n = 7/7*; EDL-Control, *n = 6/6*; EDL-PAK1 KO, *n = 7/7*. Statistics were evaluated with two two-way RM measures ANOVA. Main effects are indicated in the panels. Data are presented as mean ± S.E.M. or when applicable mean ± S.E.M. with individual data points shown. Paired data points are connected with a straight line.

### 3.3. PAK1 is dispensable for insulin-stimulated glucose uptake in lean or diet-induced insulin-resistant mice in vivo

Our findings on insulin-stimulated glucose uptake in isolated muscle from chow-fed PAK1 KO mice conflicted with a previous study reporting impaired glucose tolerance in PAK1 KO mice and defects in insulin-stimulated GLUT4 translocation in skeletal muscle *in vivo* [7]. Therefore, we further explored the effect of PAK1 KO on insulin-stimulated glucose uptake in skeletal muscle *in vivo*. Additionally, we fed a subgroup of PAK1 KO and control littermate mice a 60E% HFD for 21 weeks to investigate the role of PAK1 in insulin-resistant muscle. Insulin administration lowered blood glucose by 5.4±0.5 mM (−47%) in chow-fed control mice (Fig. 3A). In HFD-fed control mice, blood glucose dropped 3.0±0.9 mM (−26%) upon insulin administration (Fig. 3B). Lack of PAK1 had no impact on either basal blood glucose or whole-body insulin tolerance on either of the diets (Fig. 3A-B). Insulin increased glucose uptake in muscles from chow-fed (Gastrocnemius: 8.1-fold; Quadriceps: 8.5-fold, Triceps brachii: 12.3-fold; Soleus: 8.9-fold) and HFD-fed (Gastrocnemius: 3.5-fold; Quadriceps: 1.8-fold, Triceps brachii: 4.3-fold; Soleus: 11.6-fold) control mice (Fig. 3C-D). Consistent with our findings in isolated muscle, we observed no effect of PAK1 KO on basal or insulin-stimulated glucose uptake *in vivo* in muscle of either chow-fed mice or in insulin-resistant muscles from HFD-fed mice. Like glucose uptake, 2DG clearance from the plasma was unaffected by PAK1 KO in all of the investigated muscles (Fig. S1A-B). Importantly, circulating [^3^H]-2DG availability was unaffected by genotype on both of the diets (Fig. S1C). As in chow-fed mice, lack of PAK1 in HFD-fed mice had no effect on fat mass, lean body mass, body weight or energy intake (Fig. S1D-E). Similarly, whole-body glucose tolerance (Fig. S1F-G), plasma insulin concentration during the GTT (Fig. S1H) and HOMA-IR (Fig. S1I) were unaffected by PAK1 KO in HFD-fed mice. Thus, PAK1 is dispensable for *in vivo* insulin-stimulated muscle glucose uptake in both the healthy lean and the diet-induced insulin-resistant state.

**Figure 3:**
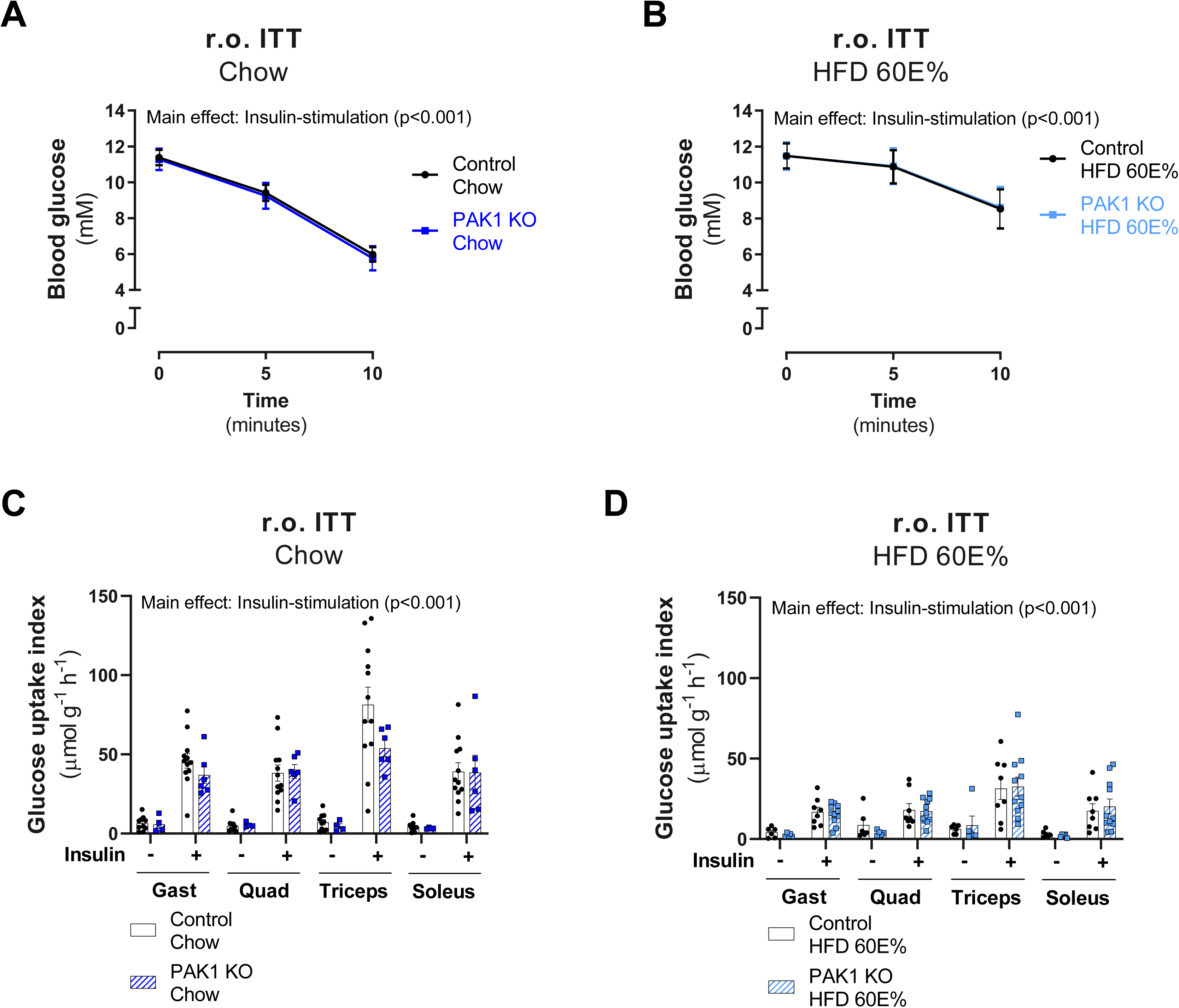
PAK1 is dispensable for insulin-stimulated glucose uptake in lean or diet-induced insulin-resistant mice in vivo. **(A-B)** Blood glucose levels during a retro-orbital insulin tolerance test (r.o. ITT) in chow- (A) and HFD-fed (B) PAK1 knockout (KO) mice or control littermates. Chow-fed mice were investigated at 10-24 weeks of age. Mice fed a 60E% high-fat diet (HFD) for 21 weeks were investigated at 27-37 weeks of age. The number of mice in each group: Chow, *n = 12/6* (Control/PAK1 KO); HFD, *n = 10/11*. Statistics were evaluated with a two-way repeated measures ANOVA. **(C-D)** Insulin-stimulated (Chow: 0.5 U kg^−1^ body weight; HFD: 60% of insulin administered to chow-fed mice) glucose uptake index in gastrocnemius (Gast), quadriceps (Quad), triceps brachii (Triceps) and soleus muscle from chow- (C) and HFD-fed (D) PAK1 KO mice or control littermates. The number of mice in each group: Chow-Saline, *n = 12/4* (Control/PAK1 KO); Chow-Insulin, *n = 12/6*; HFD-Saline, *n = 6/6*; HFD-Insulin, *n = 10/11*. Statistics were evaluated with a two-way ANOVA for each of the muscles. Main effects are indicated in the panels. Data are presented as mean ± S.E.M. or when applicable mean ± S.E.M. with individual data points shown.

### 3.4. Whole-body substrate utilization is unaffected by genetic ablation of PAK1 and PAK2

Because of discrepancies in the data resulting from the use of a global pharmacological inhibitor of group I PAKs and data resulting from PAK1 KO mice, we next sought to determine the relative contribution and involvement of PAK1 and PAK2 in insulin signalling and glucose uptake in skeletal muscle. Double knockout mice with whole-body knockout of PAK1 and muscle-specific knockout of PAK2 (1/m2 dKO) were generated as previously described [13]. By crossing 1/m2 dKO mice with littermate controls, a cohort was generated consisting of whole-body PAK1 KO, muscle-specific PAK2 (m)KO, 1/m2 dKO and littermate control mice. While no band for PAK1 could be detected in muscles lacking PAK1, muscles lacking PAK2 displayed only a partial reduction in band intensity in the immunoblots for PAK2 (Fig. 4A). This is likely due to the fact that PAK1 KO mice are whole-body knockouts, while PAK2 mKO mice are muscle-specific and other cell types within skeletal muscle tissue could thus contribute to the signal obtained in the PAK2 immunoblots. In a previous study, PAK3 was not detectable at the protein level in 1/m2 dKO muscle [13].

**Figure 4:**
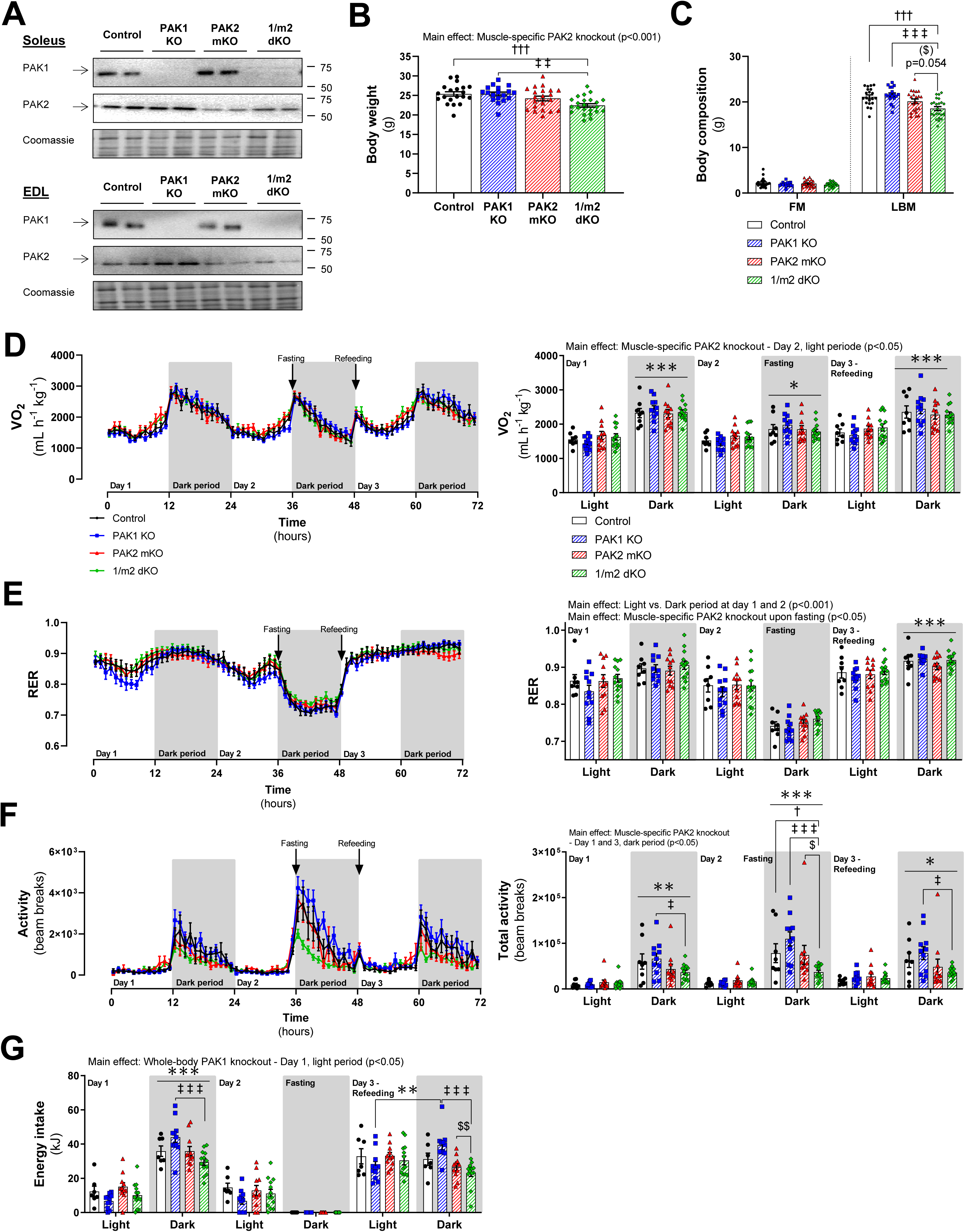
Whole-body substrate utilization is unaffected by genetic ablation of PAK1 and PAK2. **(A)** Representative blots showing PAK1 and PAK2 protein expression in soleus and extensor digitorum longus (EDL) muscle from chow-fed whole-body PAK1 knockout (KO), muscle-specific PAK2 (m)KO, PAK1/2 double KO (1/m2 dKO) mice or control littermates. **(B)** Body weight of chow-fed PAK1 KO (*n = 18*), PAK2 mKO (*n = 21*), 1/m2 dKO mice (*n = 23*) or control littermates (*n = 20*). The mice were 16-29 weeks of age. Statistics were evaluated with a two-way ANOVA to test the factors ‘PAK1’ (PAK1+/− vs. PAK−/−) and ‘PAK2’ (PAK2^fl/fl^;MyoD^+/+^ vs. PAK2^fl/fl^;MyoD^iCre/+^) thereby assessing the relative contribution of PAK1 and PAK2, respectively. Differences between genotypes were evaluated with one-way ANOVA. (**C)** Body composition (FM: Fat mass; LBM: Lean body mass) in gram in chow-fed PAK1 KO (*n = 18*), PAK2 mKO (*n = 21*), 1/m2 dKO mice (*n = 23*) or control littermates (*n = 20*). Statistics were evaluated with a two-way ANOVA to test the factors ‘PAK1’ (PAK1+/− vs. PAK−/−) and ‘PAK2’ (PAK2^fl/fl^;MyoD^+/+^ vs. PAK2^fl/fl^;MyoD^iCre/+^) thereby assessing the relative contribution of PAK1 and PAK2, respectively. Differences between genotypes were evaluated with a one-way ANOVA. **(D-G)** Oxygen uptake (VO_2_; D), respiratory exchange ratio (RER; E), activity (beam breaks; F), and energy intake (G), in chow-fed PAK1 KO (*n = 11*), PAK2 mKO (*n = 11*), 1/m2 dKO mice (*n = 13*) or control littermates (*n = 8;* for energy intake, *n = 7*) during the light and dark period recorded over a period of 72 hours. On day 2, the mice were fasted during the dark period and then refed on day 3. The mice were 23-33 weeks of age. Statistics were evaluated with two two-way ANOVAs to test the factors ‘PAK1’ (PAK1+/− vs. PAK−/−) and ‘PAK2’ (PAK2^fl/fl^;MyoD^+/+^ vs. PAK2^fl/fl^;MyoD^iCre/+^) in the light and dark period of day 1, respectively. Statistics for day 2 and 3 were evaluated similarly thereby assessing the relative contribution of PAK1 and PAK2. Differences between genotypes and the effect of the light and dark period were assessed with three two-way repeated measures (RM) ANOVA to test the factors ‘Genotype’ (Control vs. PAK1 KO vs. PAK2 mKO vs. 1/m2 dKO) and ‘Period’ (Light vs. Dark) at day 1, 2 and 3, respectively. Main effects are indicated in the panels. Significant one-way ANOVA and interactions in two-way (RM when applicable) ANOVA were evaluated by Tukey’s post hoc test: Light vs. dark period */**/*** (p<0.05/0.01/0.001); Control vs. 1/m2 dKO †/††† (p<0.05/0.001); PAK1 KO vs. 1/m2 dKO ‡/‡‡/‡‡‡ (p<0.05/0.01/0.001); PAK2 mKO vs. 1/m2 dKO ($)/$/$$ (p<0.1/0.05/0.01). Data are presented as mean ± S.E.M. or when applicable mean ± S.E.M. with individual data points shown.

As previously shown [13, 45], 1/m2 dKO mice weighed less (−12%) than control littermates (Fig. 4B, Fig. S2A-B) due to reduced (−12%) lean body mass (Fig. 4C, Fig. S2C-D). Body weight and lean body mass decreased to the same extent in 1/m2 dKO mice, leaving lean body mass percentage largely unaffected (Fig. S2E-G). Using calorimetric chambers, we monitored whole-body metabolism for 72 hours during the light and dark period. On day 2, the mice fasted during the dark period followed by refeeding on day 3. Oxygen uptake (VO_2_; Fig. 4D, Fig. S3A-B) and RER indicative of substrate utilization (Fig. 4E, Fig. S3C-D) were largely unaffected by genotype with only a slightly higher RER in mice lacking PAK2 (PAK2 mKO and 1/m2 dKO mice) upon fasting. Similar substrate utilization was obtained despite reduced habitual activity in the dark period in mice lacking PAK2, an effect largely driven by a decreased activity in 1/m2 dKO mice and increased activity in male PAK1 KO mice (Fig. 4F; Fig. S3E-F). Supporting the lower activity levels, energy intake was decreased (−11%) in 1/m2 dKO mice compared to PAK1 KO mice on day 1 (Fig. S3G) due to lower energy intake in the dark period (Fig. 4G). Upon refeeding, energy intake was reduced in mice lacking PAK2 (Fig. 4G; Fig. S3G) driven by a lower energy intake in the dark period in female mice lacking PAK2 (Fig. S3H-K). Altogether, these data suggest that lack of PAK1 and/or PAK2 are not compromising metabolic regulation during the light/dark period or in response to fasting/refeeding.

### 3.5. Glucose tolerance is reduced in mice lacking PAK2 in skeletal muscle

To test dependency on PAK1 and/or PAK2 in glucose handling and insulin sensitivity, we next investigated glucose and insulin tolerance in the 1/m2 dKO mouse strain. Blood glucose concentration in the fed state was similar between the genotypes (Fig. 5A, Fig. S4A-B). The blood glucose response to a glucose load was slightly increased in mice lacking PAK2 in skeletal muscle as evident by the increased area under the blood glucose curve (Fig. 5B-C). This was primarily driven by impaired glucose tolerance in female PAK2 mKO mice (Fig. S4C-F). Plasma insulin concentration during the GTT was unaffected by lack of PAK1 (Fig. 5D) In contrast, plasma insulin in male PAK2 mKO mice was slightly elevated compared to 1/m2 dKO mice and tended (p=0.055) to be higher than control littermates (Fig. S4G-H). This indicates impaired insulin sensitivity in male PAK2 mKO mice, but HOMA-IR was unaffected by lack of PAK1 and/or PAK2 (Fig. 5E, Fig. S5A-B). In addition, even though fasted blood glucose immediately before an ITT was modestly reduced in PAK2 mKO mice (Fig. S5C-E), the blood glucose response to an ITT was largely unaffected by lack of either PAK1 and/or PAK2 (Fig. 5F, Fig. S5F-G). Thus, despite slightly impaired glucose tolerance in mice lacking PAK2 in skeletal muscle, neither adverse effects on plasma insulin nor defects in insulin sensitivity could be detected.

### 3.6. Insulin-stimulated glucose uptake relies partially on PAK2 in EDL, but not soleus muscle, while PAK1 is not involved

To determine the relative contribution and involvement of PAK1 and PAK2 in glucose uptake in skeletal muscle, we next investigated insulin-stimulated 2DG uptake in isolated soleus and EDL. To our surprise, soleus muscle lacking PAK1 and PAK2 displayed normal insulin-stimulated 2DG uptake compared to control littermates (Fig. 6A,C). In contrast, lack of PAK2 in EDL muscle caused a modest reduction (PAK2 mKO: −18%; 1/m2 dKO: −12%) in insulin-stimulated 2DG uptake (Fig. 6B,D). Thus, in oxidative soleus muscle, group I PAKs are surprisingly dispensable for normal insulin-stimulated glucose uptake, whereas in glycolytic EDL muscle PAK2 plays a minor role.

**Figure 5:**
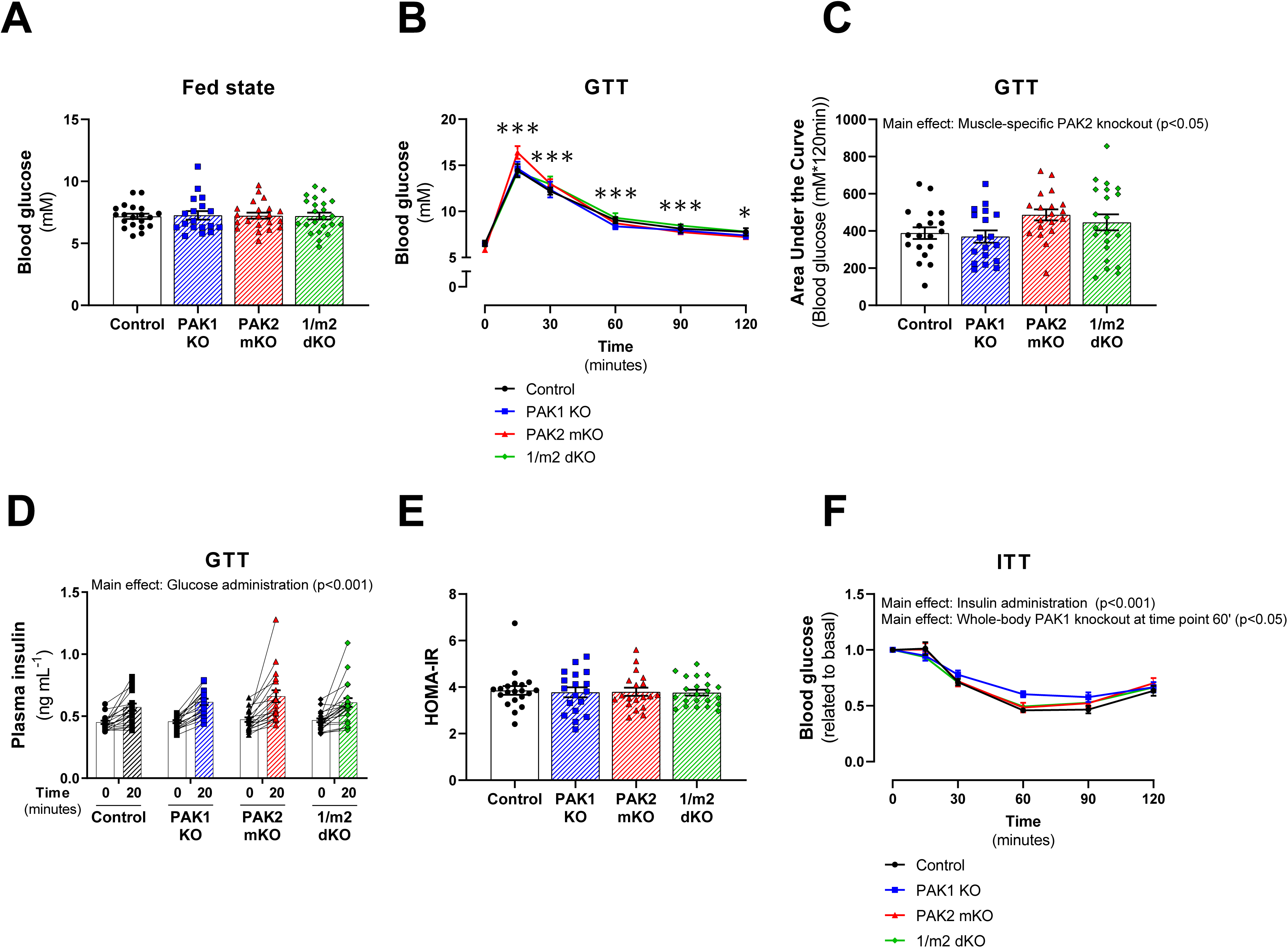
Glucose tolerance is reduced in mice lacking PAK2 in skeletal muscle. **(A)** Blood glucose concentration in the fed state (8 a.m.) in chow-fed whole-body PAK1 knockout (KO) (*n = 18*), muscle-specific PAK2 (m)KO (*n = 21*), PAK1/2 double KO (1/m2 dKO) mice (*n = 23*) or control littermates (*n = 20*). The mice were 17-30 weeks of age. Statistics were evaluated with a two-way ANOVA to test the factors ‘PAK1’ (PAK1+/− vs. PAK−/−) and ‘PAK2’ (PAK2^fl/fl^;MyoD^+/+^ vs. PAK2^fl/fl^;MyoD^iCre/+^) thereby assessing the relative contribution of PAK1 and PAK2, respectively. Differences between genotypes were evaluated with a one-way ANOVA. **(B)** Blood glucose levels during a glucose tolerance test (GTT) in chow-fed PAK1 KO (*n = 18*), PAK2 mKO (*n = 19*), 1/m2 dKO mice (*n = 21*) or control littermates (*n = 19*). The mice were 13-26 weeks of age. Statistics were evaluated with six two-way ANOVAs to test the factors ‘PAK1’ (PAK1+/− vs. PAK−/−) and ‘PAK2’ (PAK2^fl/fl^;MyoD^+/+^ vs. PAK2^fl/fl^;MyoD^iCre/+^) at time point 0, 15, 30, 60, 90 and 120, respectively, thereby assessing the relative contribution of PAK1 and PAK2. Differences between genotypes and the effect of glucose administration were assessed with a two-way repeated measures (RM) ANOVA to test the factors ‘Genotype’ (Control vs. PAK1 KO vs. PAK2 mKO vs. 1/m2 dKO) and ‘Time’ (0 vs. 15 vs. 30 vs. 60 vs. 90 vs. 120). **(C)** Incremental Area Under the Curve (AUC) for blood glucose levels during the GTT in panel B. Statistics were evaluated with a two-way ANOVA to test the factors ‘PAK1’ (PAK1+/− vs. PAK−/−) and ‘PAK2’ (PAK2^fl/fl^;MyoD^+/+^ vs. PAK2^fl/fl^;MyoD^iCre/+^) thereby assessing the relative contribution of PAK1 and PAK2, respectively. Differences between genotypes were evaluated with one-way ANOVA. **(D)** Plasma insulin values during a GTT in chow-fed PAK1 KO (*n = 17*), PAK2 mKO (*n = 19*), 1/m2 dKO mice (*n = 22*) or control littermates (*n = 19*). The mice were 15-28 weeks of age. Statistics were evaluated with two two-way ANOVAs to test the factors ‘PAK1’ (PAK1+/− vs. PAK−/−) and ‘PAK2’ (PAK2^fl/fl^;MyoD^+/+^ vs. PAK2^fl/fl^;MyoD^iCre/+^) at time point 0 and 20, respectively, thereby assessing the relative contribution of PAK1 and PAK2. Differences between genotypes and the effect of glucose administration were assessed with a two-way RM ANOVA to test the factors ‘Genotype’ (Control vs. PAK1 KO vs. PAK2 mKO vs. 1/m2 dKO) and ‘Time’ (0 vs. 20). **(E)** Homeostatic Model Assessment of Insulin Resistance (HOMA-IR) in chow-fed PAK1 KO (*n = 18*), PAK2 mKO (*n = 19*), 1/m2 dKO mice (*n = 22*) or control littermates (*n = 20*). Statistics were evaluated with a two-way ANOVA to test the factors ‘PAK1’ (PAK1+/− vs. PAK−/−) and ‘PAK2’ (PAK2^fl/fl^;MyoD^+/+^ vs. PAK2^fl/fl^;MyoD^iCre/+^) thereby assessing the relative contribution of PAK1 and PAK2, respectively. Differences between genotypes were evaluated with a one-way ANOVA. **(F)** Blood glucose levels related to the basal concentration during an insulin tolerance test (ITT) in chow-fed PAK1 KO (*n = 15*), PAK2 mKO (*n = 18*), 1/m2 dKO mice (*n = 21*) or control littermates (*n = 19*). The mice were 11-24 weeks of age. For two female control mice and four female PAK2 mKO mice, the ITT had to be stopped before the 120’-time point due to hypoglycemia (blood glucose <1.2 mM), and thus blood glucose was not determined for these mice for the last couple of time points. Statistics were evaluated with five two-way ANOVAs to test the factors ‘PAK1’ (PAK1+/− vs. PAK−/−) and ‘PAK2’ (PAK2^fl/fl^;MyoD^+/+^ vs. PAK2^fl/fl^;MyoD^iCre/+^) at time point 15, 30, 60, 90 and 120, respectively, thereby assessing the relative contribution of PAK1 and PAK2. Differences between genotypes and the effect of insulin administration were assessed with a mixed-effects model analysis to test the factors ‘Genotype’ (Control vs. PAK1 KO vs. PAK2 mKO vs. 1/m2 dKO) and ‘Time’ (0 vs. 15 vs. 30 vs. 60 vs. 90 vs. 120). Main effects are indicated in the panels. Significant one-way ANOVA and interactions in two-way (RM when applicable) ANOVA were evaluated by Tukey’s post hoc test: Effect of glucose/insulin administration vs. time point 0’ */*** (p<0.05/0.001). Data are presented as mean ± S.E.M. or when applicable mean ± S.E.M. with individual data points shown. Paired data points are connected with a straight line.

**Figure 6:**
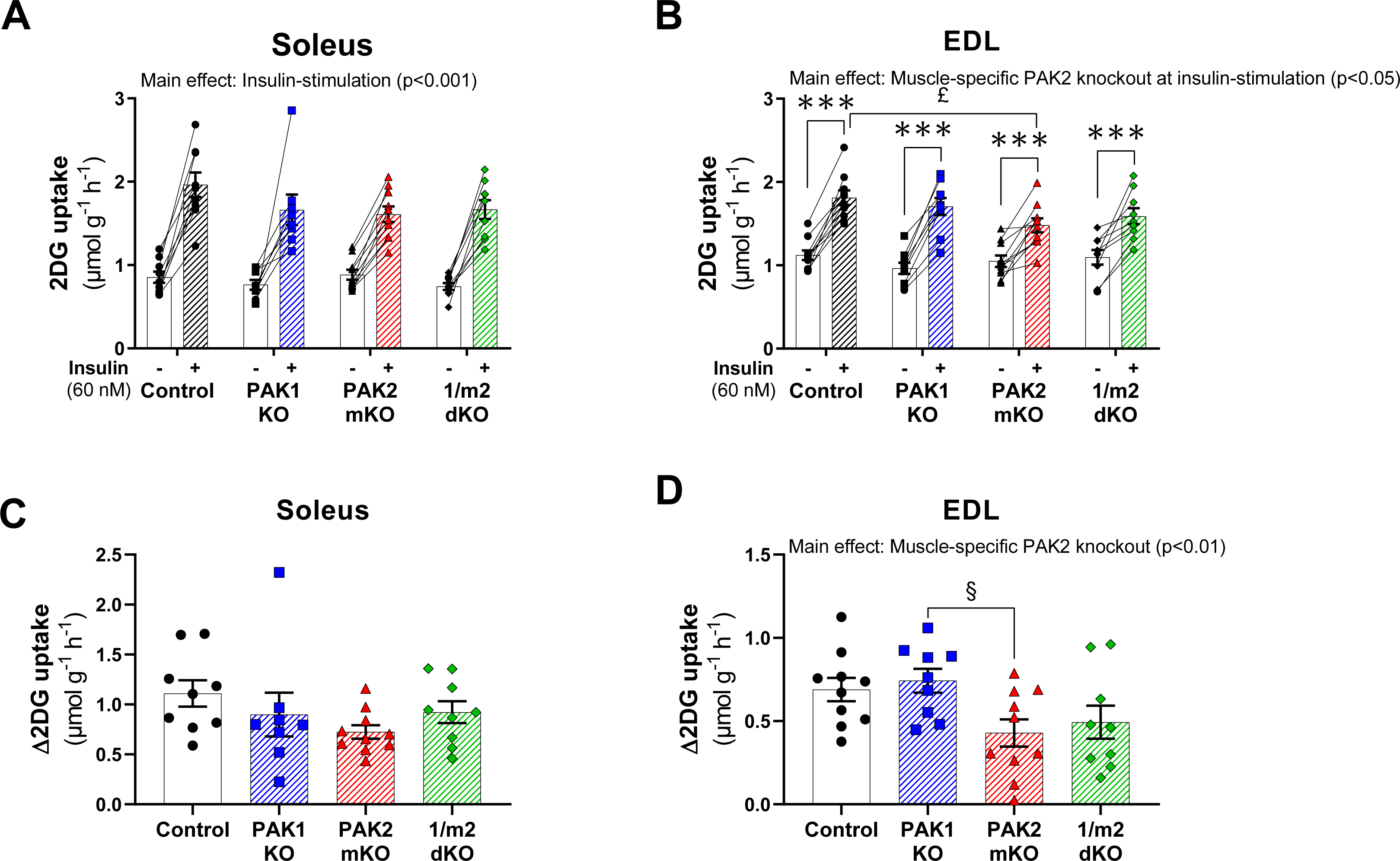
Insulin-stimulated glucose uptake relies partially on PAK2 in EDL, but not soleus muscle, while PAK1 is not involved. **(A-B)** Insulin-stimulated (60 nM) 2-Deoxyglucose (2DG) uptake in isolated soleus (A) and extensor digitorum longus (EDL; B) muscle from whole-body PAK1 knockout (KO), muscle-specific PAK2 (m)KO, PAK1/2 double KO (1/m2 dKO) mice or control littermates. Isolated muscles were pre-incubated for 20 minutes followed by 20 minutes of insulin stimulation with 2DG uptake analyzed for the final 10 minutes of stimulation. The mice were 10-16 weeks of age. Statistics were evaluated with two two-way ANOVAs to test the factors ‘PAK1’ (PAK1+/− vs. PAK−/−) and ‘PAK2’ (PAK2^fl/fl^;MyoD^+/+^ vs. PAK2^fl/fl^;MyoD^iCre/+^) in basal and insulin-stimulated samples, respectively, thereby assessing the relative contribution of PAK1 and PAK2. Differences between genotypes and the effect of insulin stimulation were assessed with a two-way repeated measures (RM) ANOVA to test the factors ‘Genotype’ (Control vs. PAK1 KO vs. PAK2 mKO vs. 1/m2 dKO) and ‘Stimuli’ (Basal vs. Insulin). **(C-D)** Δ2DG uptake in soleus (C) and EDL (D) muscle from panel C-D. Statistics were evaluated with a two-way ANOVA to test the factors ‘PAK1’ (PAK1+/− vs. PAK−/−) and ‘PAK2’ (PAK2^fl/fl^;MyoD^+/+^ vs. PAK2^fl/fl^;MyoD^iCre/+^) to assess of the relative contribution of PAK1 and PAK2, respectively. Differences between genotypes were evaluated with a one-way ANOVA. The number of determinations in each group: Control, *n = 9/10* (soleus/EDL); PAK1 KO, *n = 8/9*; PAK2 KO, *n = 10/10*; 1/m2 dKO, *n = 9/9*. Main effects are indicated in the panels. Significant one-way ANOVA and interactions in two-way (RM when applicable) ANOVA were evaluated by Tukey’s post hoc test: Insulin-stimulation vs. basal control *** (p<0.001); Control vs. PAK2 mKO £ (p<0.05); PAK1 KO vs. PAK2 mKO § (p<0.05). Data are presented as mean ± S.E.M. with individual data points shown. Paired data points are connected with a straight line.

### 3.7. PAK2 regulates TBC1D4 protein expression and signalling

Lastly, we looked into the effects of PAK1 and PAK2 on insulin-stimulated signalling. All groups displayed normal basal and insulin-stimulated pAkt S473 (Fig. 7A-B, Fig. S6A-B) and pAkt T308 (Fig. 7C-D, Fig. S6C-D) and Akt2 protein expression (Fig. S6E-F) compared to control littermates in both soleus and EDL muscle. In contrast, lack of PAK2 increased protein expression of Akt’s downstream target, TBC1D4 in soleus muscle (PAK2 mKO: +47%; 1/m2 dKO: +20%) (Fig. 7E), while reducing TBC1D4 expression in EDL (PAK2 mKO: −33%; 1/m2 dKO: −9%) (Fig. 7F). In soleus, basal and insulin-stimulated pTBC1D4 T642 was similar in all groups (Fig. 7G, Fig. S6G), suggesting that even with increased TBC1D4 expression, signalling through this pathway was normal. Concomitant with the decreased TBC1D4 expression in EDL muscle, lack of PAK2 reduced insulin-stimulated pTBC1D4 T642 (Fig. 7H, Fig. S6H) driven by attenuated (−46%) insulin-stimulated pTBC1D4 T642 in PAK2 mKO mice compared to control littermates (Fig. S6H). Knockout of TBC1D4 has been associated with lower GLUT4 protein abundance in some muscles [46, 47]. Whereas GLUT4 protein expression was normal in soleus (Fig. 7I), GLUT4 protein expression was mildly reduced in EDL in PAK2 mKO mice compared to littermate control (Fig. 7J). Thus, the slightly reduced insulin-stimulated glucose uptake in EDL muscle lacking PAK2 was concomitant with downregulated TBC1D4 signalling and GLUT4 expression supporting a role for PAK2, but not PAK1, in insulin-stimulated glucose uptake.

**Figure 7:**
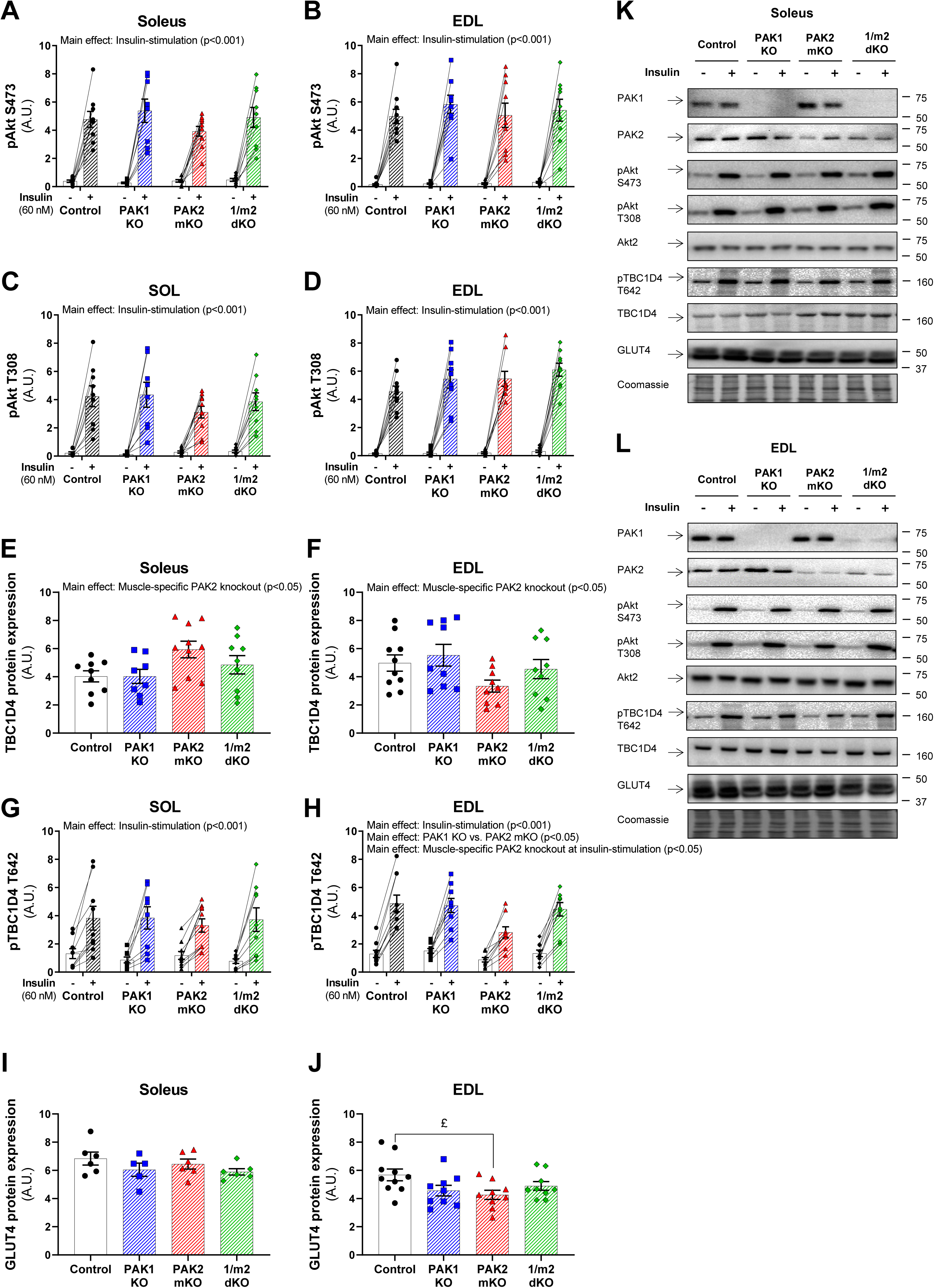
PAK2 regulates TBC1D4 protein expression and signalling. **(A-J)** Quantification of phosphorylated (p)Akt S473, pAkt T308, pTBC1D4 T642 and total TBC1D4 and GLUT4 protein expression in insulin-stimulated (60 nM) soleus (A, C, E, G, and I) and extensor digitorum longus (EDL; B, D, F, H, and J) muscle from whole-body PAK1 knockout (KO), muscle-specific PAK2 (m)KO, PAK1/2 double KO (1/m2 dKO) mice or control littermates. The mice were 10-16 weeks of age. Total protein expression is an average of the paired basal and insulin-stimulated sample. Some of the data points were excluded due to the quality of the immunoblot, so the number of determinations for GLUT4 in soleus muscle: Control, *n = 6*; PAK1 KO, *n = 5*; PAK2 KO, *n = 6*; 1/m2 dKO, *n = 6*. **(K-L)** Representative blots showing pAkt S473, pAkt T308, pTBC1D4 T642 and total PAK1, PAK2, Akt2, TBC1D4 and GLUT4 protein expression and coomassie staining as a loading control in soleus (K) and EDL (L) muscle. Statistics for phosphorylated proteins were evaluated with a two two-way ANOVAs to test the factors ‘PAK1’ (PAK1+/− vs. PAK−/−) and ‘PAK2’ (PAK2^fl/fl^;MyoD^+/+^ vs. PAK2^fl/fl^;MyoD^iCre/+^) in basal and insulin-stimulated samples, respectively, thereby assessing the relative contribution of PAK1 and PAK2. Differences between genotypes and the effect of insulin stimulation were assessed with a two-way repeated measures (RM) ANOVA to test the factors ‘Genotype’ (Control vs. PAK1 KO vs. PAK2 mKO vs. 1/m2 dKO) and ‘Stimuli’ (Basal vs. Insulin). Statistics for total protein expression were evaluated with a two-way ANOVA to test the factors ‘PAK1’ (PAK1+/− vs. PAK−/−) and ‘PAK2’ (PAK2^fl/fl^;MyoD^+/+^ vs. PAK2^fl/fl^;MyoD^iCre/+^) thereby assessing the relative contribution of PAK1 and PAK2, respectively. Differences between genotypes were evaluated with a one-way ANOVA. Main effects are indicated in the panels. Significant one-way ANOVA and interactions in two-way (RM when applicable) ANOVA were evaluated by Tukey’s post hoc test: Control vs. PAK2 mKO £ (p<0.05). Unless otherwise stated previously in the figure legend, the number of determinations in each group: Control, *n = 9/10* (soleus/EDL); PAK1 KO, *n = 8/9*; PAK2 KO, *n = 10/9*; 1/m2 dKO, *n = 9/9*. Data are presented as mean ± S.E.M. with individual data points shown. Paired data points are connected with a straight line. A.U., arbitrary units.

## 4. Discussion

We here undertook a systematic investigation into the requirement of the group I PAK isoforms in muscle glucose uptake and whole-body metabolic regulation. In contrast to previous literature, our results firmly show that PAK1 surprisingly is dispensable for insulin-stimulated glucose uptake in skeletal muscle, while PAK2 may play a minor role. Using a cohort of whole-body PAK1 KO mice and another cohort of transgenic mice lacking either PAK1 (whole-body), PAK2 (muscle-specific), or jointly lacking both PAK1 and muscle PAK2, we show that PAK1 is not required in insulin-stimulated muscle glucose uptake *in vivo* or in isolated muscles. In accordance, we found no effect of whole-body PAK1 KO on glucose tolerance in either mice fed a standard chow diet (insulin sensitive mice) or in mice fed a HFD (insulin-resistant mice). In contrast, PAK2 seemed partially involved in insulin-stimulated glucose uptake, in glycolytic EDL muscle. This could potentially explain the slightly impaired glucose tolerance with the muscle-specific knockout of PAK2 in mice. Nevertheless, supporting only a minor role for skeletal muscle PAK2 in the whole-body substrate utilization, RER was largely unaffected by lack of PAK1 and/or PAK2 and only slightly elevated in mice lacking PAK2 when challenged by fasting.

In a previous study, the increase in GLUT4 abundance at the plasma membrane in response to insulin measured was completely abrogated in PAK1 KO muscle as measured by subcellular fractionation of homogenates of hindlimb skeletal muscles [7], suggesting that PAK1 is necessary for insulin-stimulated GLUT4 translocation. Although glucose uptake was not assessed in that study [7], this indicated a key role for PAK1 in regulating glucose uptake in mouse skeletal muscle. Surprisingly, PAK1 was not required for insulin-stimulated glucose uptake in our hands. Furthermore, in our study, both chow- and HFD-fed PAK1 KO mice displayed blood glucose concentrations similar to control littermates during a GTT. This was in contrast to previous studies reporting impaired glucose tolerance in chow-fed PAK1 KO mice [7, 27] and elevated fasting blood glucose in PAK1 KO mice fed a western diet (45E% fat) [44]. Instead, despite the previous finding that insulin-stimulated GLUT4 translocation was unaffected by a 75% knockdown of PAK2 in L6-GLUT4myc myoblasts [6], we found that muscle-specific PAK2 KO slightly impaired glucose tolerance and insulin-stimulated glucose in mouse skeletal muscle. These discrepancies between our and previous findings are difficult to delineate but might be due to methodological differences. Wang et al. [7] used a crude fractionation method to measure GLUT4 translocation, whereas we analyzed the direct outcome hereof: glucose uptake. Although the insulin-induced increase in 3-O-methylglucose transport correlates with the increase in cell surface GLUT4 protein content in human skeletal muscle strips [48], discrepancies between the presence of GLUT4 at the plasma membrane and glucose uptake have occasionally been reported in cell culture studies [49–51]. Another potential explanatory factor could be that our studies were conducted in both female and male mice, whereas past studies in PAK1 KO mice have been conducted in 4-6 months old male mice [7, 44]. However, our data suggest no major differences between female and male mice in the response to lack of PAK1 and/or PAK2 on the whole-body metabolic parameters measured. Instead, the discrepancies between our and previous findings could be due to an effect of age, as our studies were conducted in mice at different ranges of age (10-37 of weeks age at the terminal experiment). In fact, age-dependent myopathy and development of megaconial mitochondria have been reported in the 1/m2 dKO mice [45]. Regardless, even though a role for group I PAKs in age-related insulin resistance should be further investigated, our investigation suggests that group I PAKs are largely dispensable in regulating whole-body glucose homeostasis or insulin-stimulated glucose uptake in skeletal muscle.

In the current study, pharmacological inhibition of group I PAKs inhibited muscle glucose uptake in response to insulin. Similarly, IPA-3 previously inhibited insulin-stimulated GLUT4 translocation and glucose uptake in L6-GLUT4myc myoblasts and myotubes, respectively [6]. IPA-3 is a non-ATP-competitive allosteric inhibitor of all group I PAKs (PAK1, 2, and 3). IPA-3 binds covalently to the regulatory CRIB domain of group I PAKs, thereby preventing binding to PAK activators, such as Rac1 [43]. Although IPA-3 is reported to be a highly selective and well-described inhibitor of group I PAKs that does not affect other groups of PAKs or similar kinases tested [43], pharmacological inhibitors often have off-target effects [52] which is a concern. It is also possible that acute IPA-3-induced inhibition of group I PAKs elicits more potent effects compared with jointly knockout of PAK1 and PAK2 because the transgenic manipulations have been present from birth and may thus have resulted in compensatory changes. The development of inducible muscle-specific group I PAK deficient models could help clarify this. Importantly, any possible compensatory mechanisms cannot be via redundancy with group I PAKs, as PAK1 and PAK2 are removed genetically, and even in 1/m2 dKO mice, PAK3 cannot be detected at the protein level [13]. This emphasizes that group I PAKs are largely dispensable for insulin-stimulated glucose uptake in skeletal muscle with only PAK2 playing a minor role.

Our hypothesis was that group I PAKs would be significantly involved in insulin-stimulated glucose uptake because of the established necessity of their upstream activator, Rac1 [15–20]. Our findings suggest that Rac1 does not exclusively mediate insulin-stimulated glucose uptake through group I PAKs. Another downstream target of Rac1 is RalA. GLUT4 translocation induced by a constitutively activated Rac1 mutant was abrogated in L6-GLUT4myc myoblasts upon RalA knockdown [53] and, importantly, overexpression of a dominant-negative mutant of RalA reduced GLUT4 translocation in response to insulin in mouse gastrocnemius muscle fibres [54]. Additionally, Rac1 is an essential component in the activation of the NADPH oxidase (NOX) complex [55, 56]. In L6-GLUT4myc myotubes, reactive oxygen species have been reported to induce NOX2-dependent GLUT4 translocation in response to insulin [57]. Insulin-stimulated NOX2 regulation in mature muscle remains to be investigated. However, a recent study suggested a role for Rac1 in the regulation of muscle glucose uptake through activation of the NOX2 in response to exercise [58]. Since Rac1 is required for both contraction- and insulin-stimulated glucose uptake in isolated mouse muscle [18, 59], Rac1 could also be involved in insulin-stimulated glucose uptake via NOX2 activation. Consequently, future studies should aim to investigate other players downstream of Rac1 since group I PAKs seem to be largely dispensable for glucose uptake in mature skeletal muscle.

## 5. Conclusion

Based on our present findings, we conclude that PAK2 may be a requirement for insulin-stimulated glucose uptake in EDL muscle. However, in contrast to previous reports, group I PAKs are largely dispensable in the regulation of whole-body glucose homeostasis and insulin-stimulated glucose uptake in mouse skeletal muscle.

## Author contributions

**LLVM:** Conceptualization; Methodology; Formal analysis; Investigation; Writing - Original Draft; Writing - Review & Editing; Visualization; Project administration; Funding acquisition. **MJ:** Investigation; Writing - Review & Editing. **RK:** Methodology; Investigation; Writing - Review & Editing. **GAJ:** Resources; Writing - Review & Editing. **ABM:** Investigation; Writing - Review & Editing; Funding acquisition. **JRK:** Investigation; Writing - Review & Editing; Funding acquisition. **A-ML:** Investigation; Writing - Review & Editing; Funding acquisition. **NRA:** Investigation; Writing - Review & Editing. **PS:** Methodology; Investigation; Writing - Review & Editing. **TEJ:** Investigation; Writing - Review & Editing; Funding acquisition. **RSK:** Resources; Writing - Review & Editing; Funding acquisition. **EAR:** Conceptualization; Methodology; Writing - Original Draft; Writing - Review & Editing; Supervision; Project administration; Funding acquisition. **LS:** Conceptualization; Methodology; Investigation; Writing - Original Draft; Writing - Review & Editing; Supervision; Project administration; Funding acquisition. LS is the guarantor of this work and takes responsibility for the integrity of the data and the accuracy of the data analysis.

## Acknowledgements

We thank our colleagues, especially Jørgen Wojtaszewski and Bente Kiens, at the Section of Molecular Physiology, Department of Nutrition, Exercise, and Sports (NEXS), Faculty of Science, University of Copenhagen, for fruitful discussions on this topic. We acknowledge the skilled technical assistance of Betina Bolmgren, Irene B. Nielsen, and Mona Ali (Section of Molecular Physiology, NEXS, Faculty of Science, University of Copenhagen, Denmark). The PAK1 KO founder mice were a kind gift from Debbie Thurmond (Department of Molecular & Cellular Endocrinology, Diabetes and Metabolism Research Institute, City of Hope/BRICA, USA). The graphical abstract was modified from Servier Medical Art, licensed under a Creative Common Attribution 3.0 Generic License (http://smart.servier.com/).

## Grants

This study was supported by a PhD fellowship from The Lundbeck Foundation (grant 2015-3388 to LLVM); PhD scholarships from The Danish Diabetes Academy, funded by The Novo Nordisk Foundation (ABM and JRK); Postdoctoral research grant from the Danish Diabetes Academy, funded by the Novo Nordisk Foundation (grant NNF17SA0031406 to A-ML); National Institute of Arthritis and Musculoskeletal and Skin Diseases (grant AR046207 and AR070231 to RSK); The Danish Council for Independent Research, Medical Sciences (grant DFF-4004-00233 to LS, grant 6108-00203 to EAR); The Novo Nordisk Foundation (grant 10429 to EAR, grant 15182 to TEJ, grant NNF16OC0023418 and NNF18OC0032082 to LS).

## Appendix A. Supplementary data

Supplementary table S1 and supplementary figures S1-6 are to be found in Appendix A.

## Conflict of interest

None declared.

## Data and Resource Availability

The datasets generated and analyzed during the current study are available from the corresponding author upon reasonable request. No novel applicable resources were generated or analyzed during the current study.

## Appendix A. Supplementary data

**Table S1.**
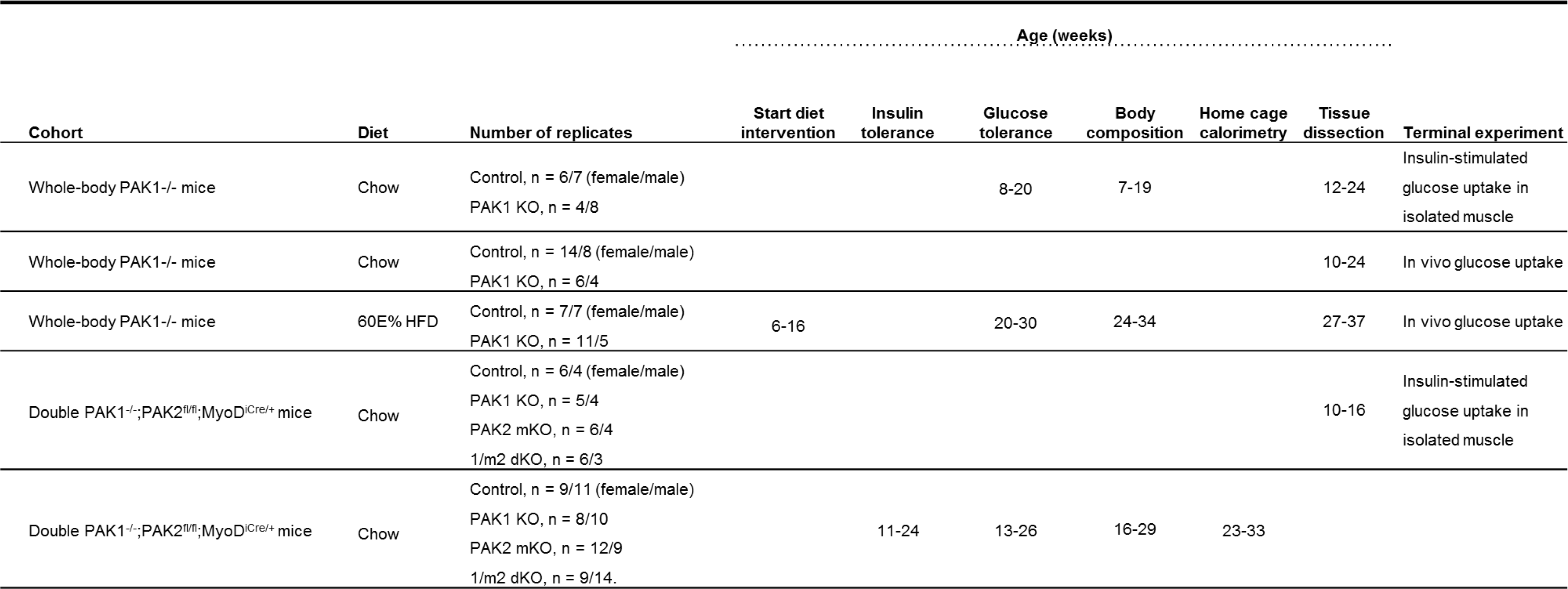
Overview of the different cohorts of genetically modified mice

**Figure S1:**
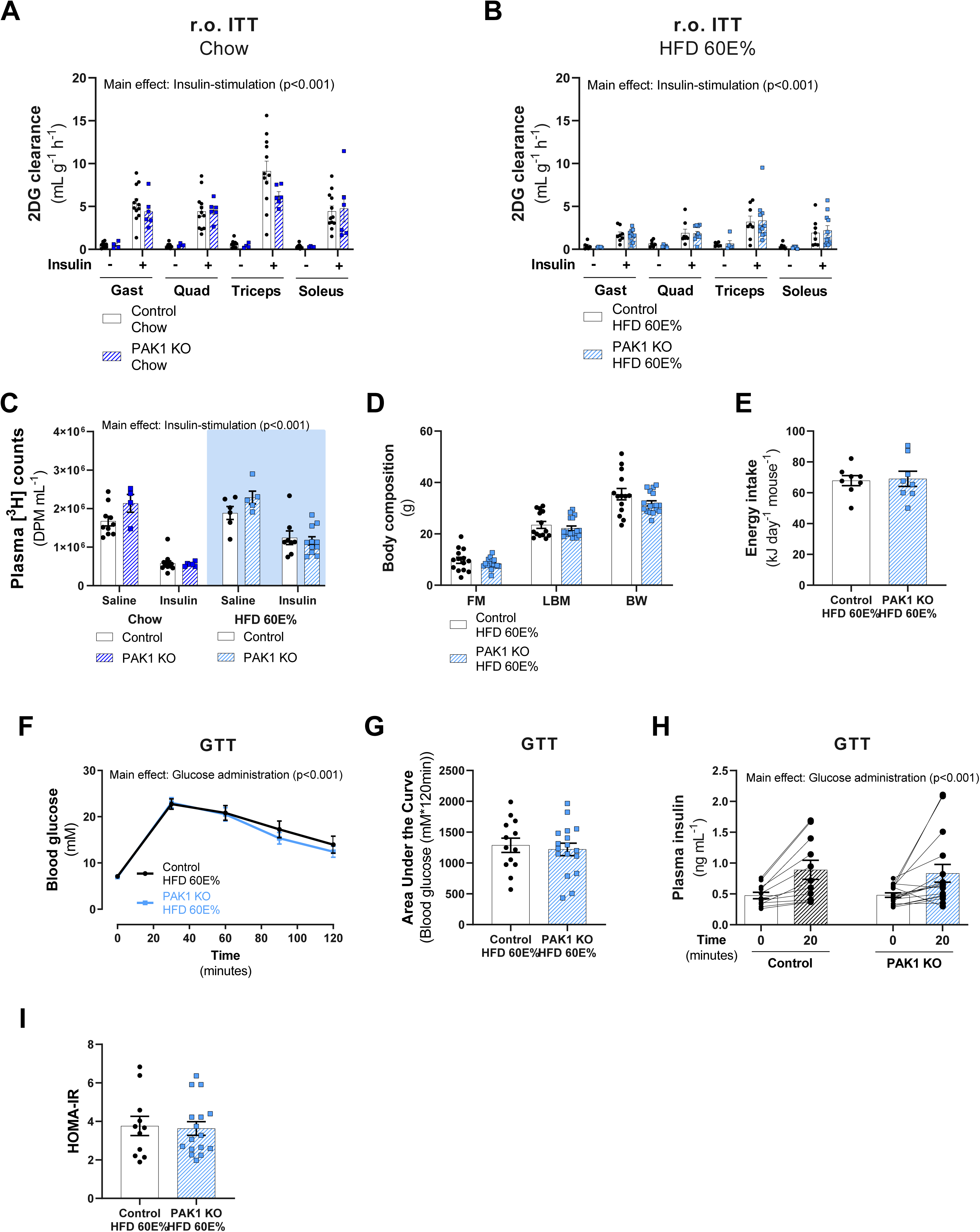
**(A-B)** Insulin-stimulated (Chow: 0.5 U kg^−1^ body weight; HFD: 60% of insulin administered to chow-fed mice) 2-Deoxyglucose (2DG) clearance in gastrocnemius (Gast), quadriceps (Quad), triceps brachii (Triceps) and soleus muscle from chow- (A) and 60E% high-fat diet (HFD)-fed (B) PAK1 knockout (KO) mice or control littermates. The number of mice in each group: Chow-Saline, *n = 12/4* (Control/PAK1 KO); Chow-Insulin, *n = 12/6*; HFD-Saline, *n = 6/6*; HFD-Insulin, *n = 10/11*. Statistics were evaluated with a two-way ANOVA for each of the muscles. **(C)** Plasma [^3^H] counts 10 minutes after retro-orbital (r.o.) administration of a bolus of saline containing [^3^H]-labelled 2DG ([^3^H]-2DG) with or without insulin. Statistics were evaluated with two two-way ANOVAs to test the factors ‘stimuli’ (basal vs. insulin) and ‘genotype’ (control vs. PAK1 KO) in chow-fed and HFD-fed mice, respectively. **(D)** Body composition (FM: Fat mass; LBM: Lean body mass; BW: Body weight) in gram in HFD-fed PAK1 KO mice (*n = 17*) or control littermates (*n = 14*). Body composition was assessed in week 18-19 of the diet intervention. The mice were 24-34 weeks of age. Statistics were evaluated with a Student’s t-test. **(E)** Energy intake in HFD-fed PAK1 KO mice (*n = 8*) or control littermates (*n = 8*). Energy intake was monitored in a separate cohort of mice. Statistics were evaluated with a Student’s t-test. **(F)** Blood glucose levels during a glucose tolerance test (GTT) in HFD-fed PAK1 KO mice (*n = 17*) or control littermates (*n = 13*). Glucose tolerance was assessed in week 14 of the diet intervention. The mice were 20-30 weeks of age. Statistics were evaluated with a two-way repeated measures (RM) ANOVA. **(G)** Incremental Area Under the Curve (AUC) for blood glucose levels during the GTT in panel F. Statistics were evaluated with a Student’s t-test. **(H)** Plasma insulin values during a GTT in HFD-fed PAK1 KO mice (*n = 16*) or control littermates (*n = 11*). The plasma insulin response to glucose administration was measured in week 16 of the diet intervention. The mice were 22-32 weeks of age. Statistics were evaluated with a two-way RM ANOVA. **(I)** Homeostatic Model Assessment of Insulin Resistance (HOMA-IR) in HFD-fed PAK1 KO mice (*n = 16*) or control littermates (*n = 11*). Statistics were evaluated with a Student’s t-test. Main effects are indicated in the panels. Data are presented as mean ± S.E.M. or when applicable mean ± S.E.M. with individual data points shown. Paired data points are connected with a straight line.

**Figure S2:**
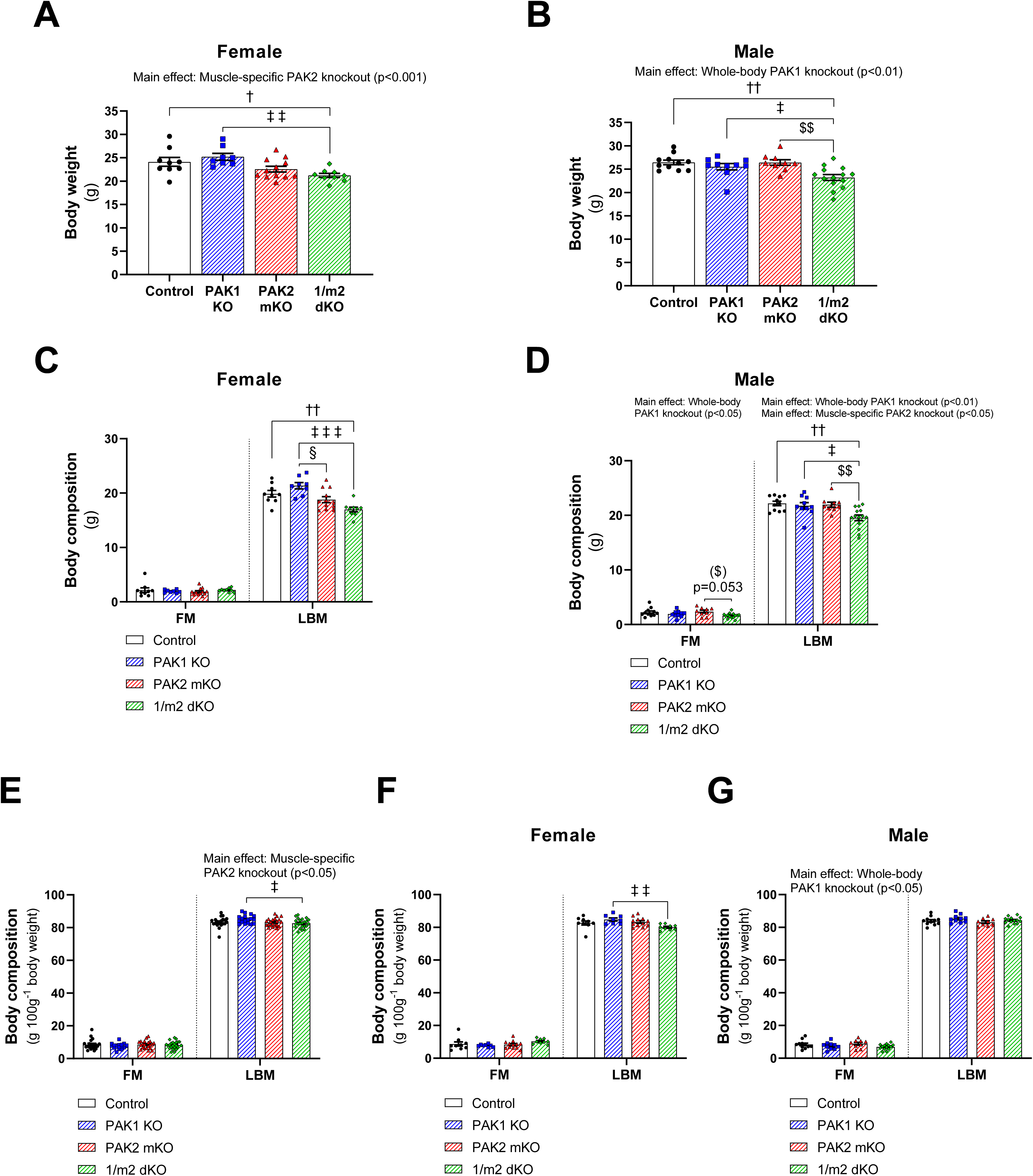
**(A-B)** Body weight of female (A) and male (B) whole-body PAK1 knockout (KO), muscle-specific PAK2 (m)KO, PAK1/2 double KO (1/m2 dKO) mice or control littermates. **(C-D)** Body composition (FM: Fat mass; LBM: Lean body mass) in gram in female (C) and male (D) PAK1 KO, PAK2 mKO, 1/m2 dKO or control littermates. **(E-G)** Body composition (FM: Fat mass; LBM: Lean body mass) in percentage in both sexes combined (E) and in female (F) and male (G) PAK1 KO, PAK2 mKO, 1/m2 dKO or control littermates. The number of mice in each group: Control, *n = 9/11* (female/male); PAK1 KO, *n = 8/10*; PAK2 mKO, *n = 12/9*; 1/m2 dKO, *n = 9/14*. Statistics were evaluated with a two-way ANOVA to test the factors ‘PAK1’ (PAK1+/− vs. PAK−/−) and ‘PAK2’ (PAK2^fl/fl^;MyoD^+/+^ vs. PAK2^fl/fl^;MyoD^iCre/+^) thereby assessing the relative contribution of PAK1 and PAK2, respectively. Differences between genotypes were evaluated with a one-way ANOVA. Main effects are indicated in the panels. Significant one-way ANOVA and interactions in two-way ANOVA were evaluated by Tukey’s post hoc test: Control vs. 1/m2 dKO †/†† (p<0.05/0.01); PAK1 KO vs. PAK2 mKO § (p<0.05); PAK1 KO vs. 1/m2 dKO ‡/‡‡/‡‡‡ (p<0.05/0.01/0.001); PAK2 mKO vs. 1/m2 dKO ($)/$$ (p<0.1/0.01). Data are presented as mean ± S.E.M. with individual data points shown.

**Figure S3:**
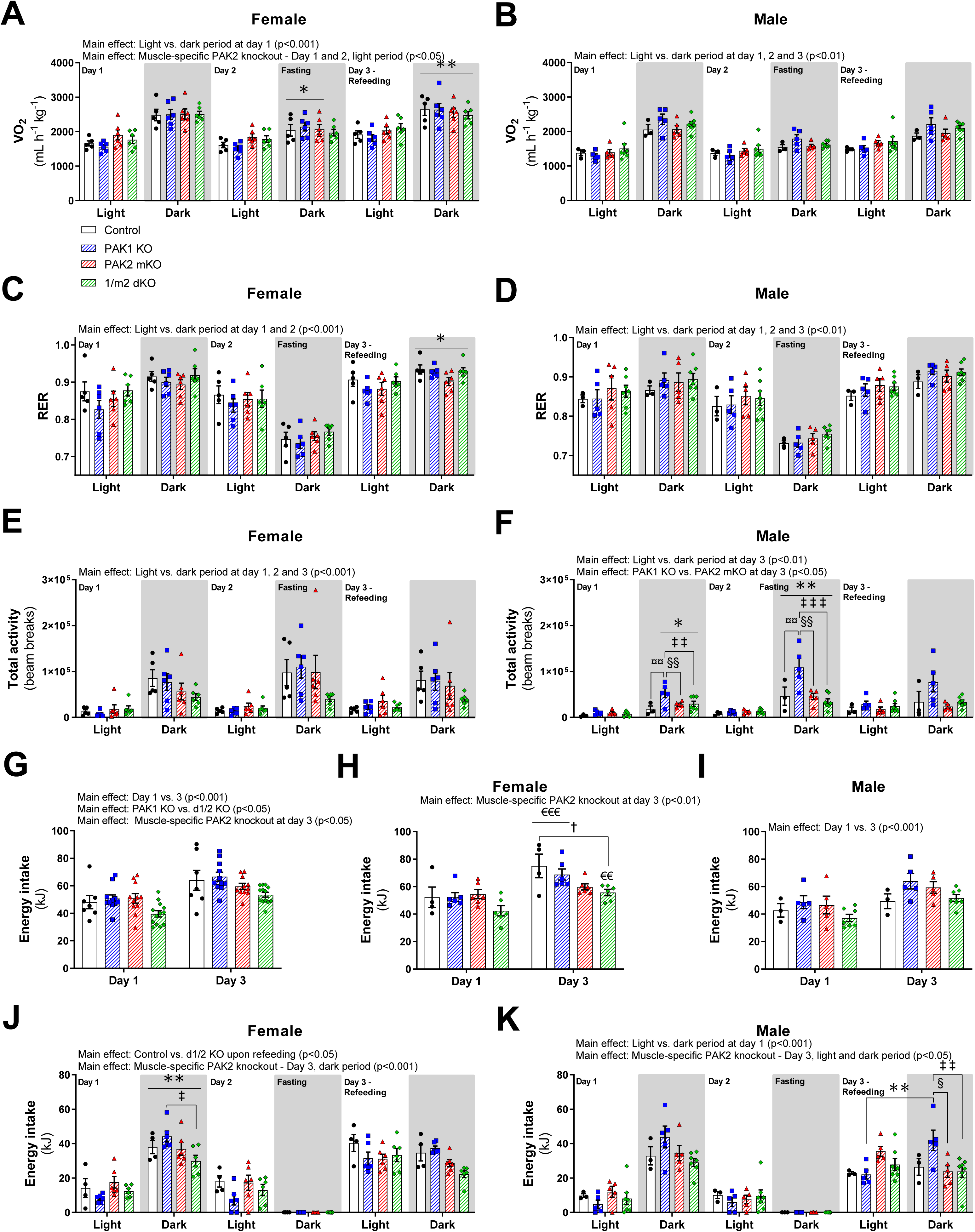
**(A-F)** Oxygen uptake (VO_2_), respiratory exchange ratio (RER), and activity (beam breaks) in female (A, C, and E) and male (B, D, and F) whole-body PAK1 knockout (KO), muscle-specific PAK2 (m)KO, PAK1/2 double KO (1/m2 dKO) mice or control littermates during the light and dark period recorded over a period of 72 hours in calorimetric chambers. On day 2, mice fasted during the dark period and then refed on day 3. **(G-I)** Total energy intake on day 1 and day 3 in both sexes combined (G) and in female (H) and male (I) whole-body PAK1 KO, PAK2 mKO, 1/m2 dKO mice or control littermates recorded over a period of 72 hours in calorimetric chambers. **(J-K)** Energy intake in female (J) and male (K) whole-body PAK1 KO, Pak2 mKO, 1/m2 dKO mice or control littermates during the light and dark period recorded over a period of 72 hours in calorimetric chambers. On day 2, mice fasted during the dark period and were then refed on day 3. The number of mice in each group: Control, *n = 5/3* (female/male; for energy intake, *n = 4/3*); PAK1 KO, *n = 6/5*; PAK2 mKO, *n = 6/5*; 1/m2 dKO, *n = 6/7*. **(A-F+J-K)** Statistics were evaluated with two two-way ANOVAs to test the factors ‘PAK1’ (PAK1+/− vs. PAK−/−) and ‘PAK2’ (PAK2^fl/fl^;MyoD^+/+^ vs. PAK2^fl/fl^;MyoD^iCre/+^) in the light and dark period of day 1, respectively. Statistics for day 2 and 3 were evaluated similarly thereby assessing the relative contribution of PAK1 and PAK2. Differences between genotypes and the effect of the light and the dark period were assessed with three two-way repeated measures (RM) ANOVA to test the factors ‘Genotype’ (Control vs. PAK1 KO vs. PAK2 mKO vs. 1/m2 dKO) and ‘Period’ (Light vs. Dark) at day 1, 2 and 3, respectively. **(G-I)** Statistics were evaluated with two two-way ANOVAs to test the factors ‘PAK1’ (PAK1+/− vs. PAK−/−) and ‘PAK2’ (PAK2^fl/fl^;MyoD^+/+^ vs. PAK2^fl/fl^;MyoD^iCre/+^) at day 1 and day 2, respectively. Differences between genotypes and the day were assessed with one two-way repeated measures (RM) ANOVA to test the factors ‘Genotype’ (Control vs. PAK1 KO vs. PAK2 mKO vs. 1/m2 dKO) and ‘Day’ (Day 1 vs. Day 3). Main effects are indicated in the panels. Significant one-way ANOVA and interactions in two-way (RM when applicable) ANOVA were evaluated by Tukey’s post hoc test: Light vs. dark period */**/*** (p<0.05/0.01/0.001); Day 1 vs. Day 3 €€/€€€ (p<0.01/0.001); Control vs. PAK1 KO ¤¤ (p<0.01); Control vs. 1/m2 dKO † (p<0.05); PAK1 KO vs. PAK2 mKO §/§§ (p<0.05/0.01); PAK1 KO vs. 1/m2 dKO ‡/‡‡/‡‡‡ (p<0.05/0.01/0.001). Data are presented as mean ± S.E.M. with individual data points shown.

**Figure S4:**
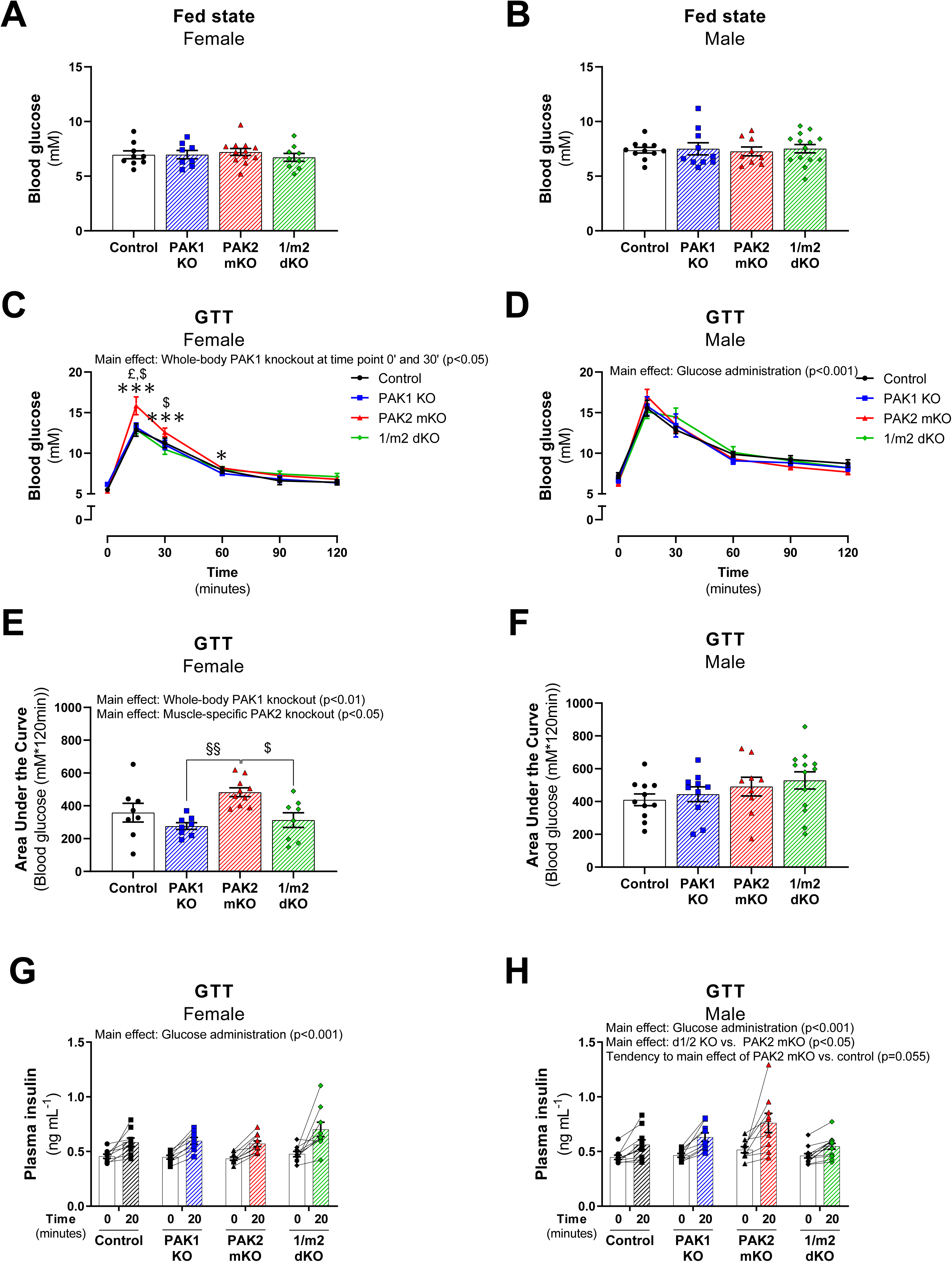
**(A-B)** Blood glucose concentration in the fed state (8 a.m.) in female (A) and male (B) whole-body PAK1 knockout (KO), muscle-specific PAK2 (m)KO, PAK1/2 double KO (1/m2 dKO) mice or control littermates. The number of mice in each group: Control, *n = 9/11* (female/male); PAK1 KO, *n = 8/10*; PAK2 mKO, *n = 12/9*; 1/m2 dKO, *n = 9/14*. Statistics were evaluated with a two-way ANOVA to test the factors ‘PAK1’ (PAK1+/− vs. PAK−/−) and ‘PAK2’ (PAK2^fl/fl^;MyoD^+/+^ vs. PAK2^fl/fl^;MyoD^iCre/+^) thereby assessing the relative contribution of PAK1 and PAK2, respectively. Differences between genotypes were evaluated with a one-way ANOVA. **(C-D)** Blood glucose levels during a glucose tolerance test (GTT) in female (C) and male (D) PAK1 KO, PAK2 mKO, 1/m2 dKO mice or control littermates. The number of mice in each group: Control, *n = 8/11* (female/male); PAK1 KO, *n = 8/10*; PAK2 mKO, *n = 10/9*; 1/m2 dKO, *n = 8/13*. Statistics were evaluated with six two-way ANOVAs to test the factors ‘PAK1’ (PAK1+/− vs. PAK−/−) and ‘PAK2’ (PAK2^fl/fl^;MyoD^+/+^ vs. PAK2^fl/fl^;MyoD^iCre/+^) at time point 0, 15, 30, 60, 90 and 120, respectively, thereby assessing the relative contribution of PAK1 and PAK2. Differences between genotypes and the effect of glucose administration were assessed with a two-way repeated measures (RM) ANOVA to test the factors ‘Genotype’ (Control vs. PAK1 KO vs. PAK2 mKO vs. 1/m2 dKO) and ‘Time’ (0 vs. 15 vs. 30 vs. 60 vs. 90 vs. 120). **(E-F)** Incremental Area Under the Curve (AUC) for blood glucose levels for females (E) and male (F) mice during the GTT in panel C-D. Statistics were evaluated with a two-way ANOVA to test the factors ‘PAK1’ (PAK1+/− vs. PAK−/−) and ‘PAK2’ (PAK2^fl/fl^;MyoD^+/+^ vs. PAK2^fl/fl^;MyoD^iCre/+^) thereby assessing the relative contribution of PAK1 and PAK2, respectively. Differences between genotypes were evaluated with a one-way ANOVA. **(G-H)** Plasma insulin values during a GTT in female (G) and male (F) PAK1 KO, PAK2 mKO, 1/m2 dKO mice or control littermates. The number of mice in each group: Control, *n = 9/10* (female/male); PAK1 KO, *n = 8/9*; PAK2 mKO, *n = 10/9*; 1/m2 dKO, *n = 9/13*. Statistics were evaluated with two two-way ANOVAs to test the factors ‘PAK1’ (PAK1+/− vs. PAK−/−) and ‘PAK2’ (PAK2^fl/fl^;MyoD^+/+^ vs. PAK2^fl/fl^;MyoD^iCre/+^) at time point 0 and 20, respectively, thereby assessing the relative contribution of PAK1 and PAK2. Differences between genotypes and the effect of glucose administration were assessed with a two-way RM ANOVA to test the factors ‘Genotype’ (Control vs. PAK1 KO vs. PAK2 mKO vs. 1/m2 dKO) and ‘Time’ (0 vs. 20). Main effects are indicated in the panels. Significant one-way ANOVA and interactions in two-way (RM when applicable) ANOVA were evaluated by Tukey’s post hoc test: Effect of glucose administration vs. time point 0’ */*** (p<0.05/0.001); Control vs. PAK2 mKO £ (p<0.05); PAK1 KO vs. PAK2 mKO §§ (p<0.01); PAK2 mKO vs. 1/m2 dKO $ (p<0.05). Data are presented as mean ± S.E.M. or when applicable mean ± S.E.M. with individual data points shown. Paired data points are connected with a straight line.

**Figure S5:**
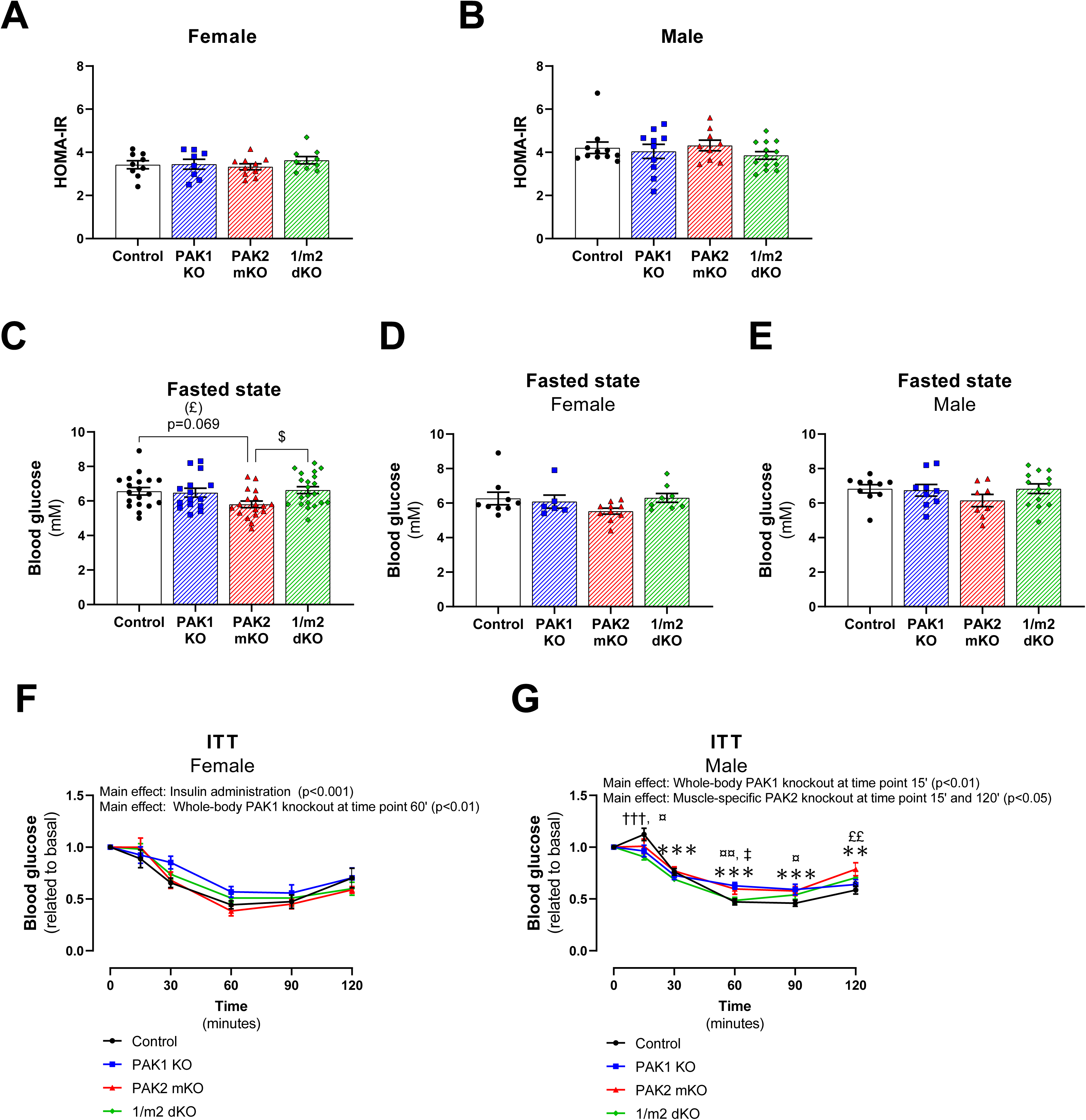
**(A-B)** Homeostatic Model Assessment of Insulin Resistance (HOMA-IR) in female (A) and male (B) whole-body PAK1 knockout (KO), muscle-specific PAK2 (m)KO, PAK1/2 double KO (1/m2 dKO) mice or control littermates. The number of mice in each group: Control, *n = 9/11* (female/male); PAK1 KO, *n = 8/10*; PAK2 mKO, *n = 10/9*; 1/m2 dKO, *n = 9/13*. Statistics were evaluated with a two-way ANOVA to test the factors ‘PAK1’ (PAK1+/− vs. PAK−/−) and ‘PAK2’ (PAK2^fl/fl^;MyoD^+/+^ vs. PAK2^fl/fl^;MyoD^iCre/+^) thereby assessing the relative contribution of PAK1 and PAK2, respectively. Genotype differences were evaluated with a one-way ANOVA. **(C-E)** Basal blood glucose concentration (fasted state) immediately before an insulin tolerance test (ITT) in both sexes combined (C) and in female (D) and male (E) PAK1 KO, PAK2 mKO, 1/m2 dKO mice and control littermates. The number of mice in each group: Control, *n = 9/10* (female/male); PAK1 KO, *n = 6/9*; PAK2 mKO, *n = 10/8*; 1/m2 dKO, *n = 8/13*. Statistics were evaluated with a two-way ANOVA to test the factors ‘PAK1’ (PAK1+/− vs. PAK−/−) and ‘PAK2’ (PAK2^fl/fl^;MyoD^+/+^ vs. PAK2^fl/fl^;MyoD^iCre/+^) thereby assessing the relative contribution of PAK1 and PAK2, respectively. Differences between genotypes were evaluated with a one-way ANOVA. **(F-G)** Blood glucose levels related to basal concentration during an ITT in female (F) and male (G) PAK1 KO, PAK2 mKO, 1/m2 dKO mice or control littermates. The number of mice in each group: Control, *n = 9/10* (female/male); PAK1 KO, *n = 6/9*; PAK2 mKO, *n = 10/8*; 1/m2 dKO, *n = 8/13*. For two female control mice and four female PAK2 mKO mice, the ITT had to be stopped before the 120’-time point due to hypoglycemia (blood glucose <1.2 mM), and thus blood glucose was not determined for these mice for the last couple of time points. Statistics were evaluated with five two-way ANOVAs to test the factors ‘PAK1’ (PAK1+/− vs. PAK−/−) and ‘PAK2’ (PAK2^fl/fl^;MyoD^+/+^ vs. PAK2^fl/fl^;MyoD^iCre/+^) at time point 15, 30, 60, 90 and 120, respectively, thereby assessing the relative contribution of PAK1 and PAK2. Differences between genotypes and the effect of insulin administration were assessed with a two-way repeated measures (RM) ANOVA (mixed-effects model analysis for female mice) to test the factors ‘Genotype’ (Control vs. PAK1 KO vs. PAK2 mKO vs. 1/m2 dKO) and ‘Time’ (0 vs. 15 vs. 30 vs. 60 vs. 90 vs. 120). Main effects are indicated in the panels. Significant one-way ANOVA and interactions in two-way (RM when applicable) ANOVA were evaluated by Tukey’s post hoc test: Effect of insulin administration vs. time point 0’ **/*** (p<0.01/0.001); Control vs. PAK1 KO ¤/¤¤ (p<0.05/0.01); Control vs. PAK2 mKO (£)££ (p<0.1/0.01); Control vs. 1/m2 dKO ††† (p<0.001); PAK1 KO vs. 1/m2 dKO ‡ (p<0.05); PAK2 mKO vs. 1/m2 dKO $ (p<0.05). Data are presented as mean ± S.E.M. or when applicable mean ± S.E.M. with individual data points shown.

**Figure S6:**
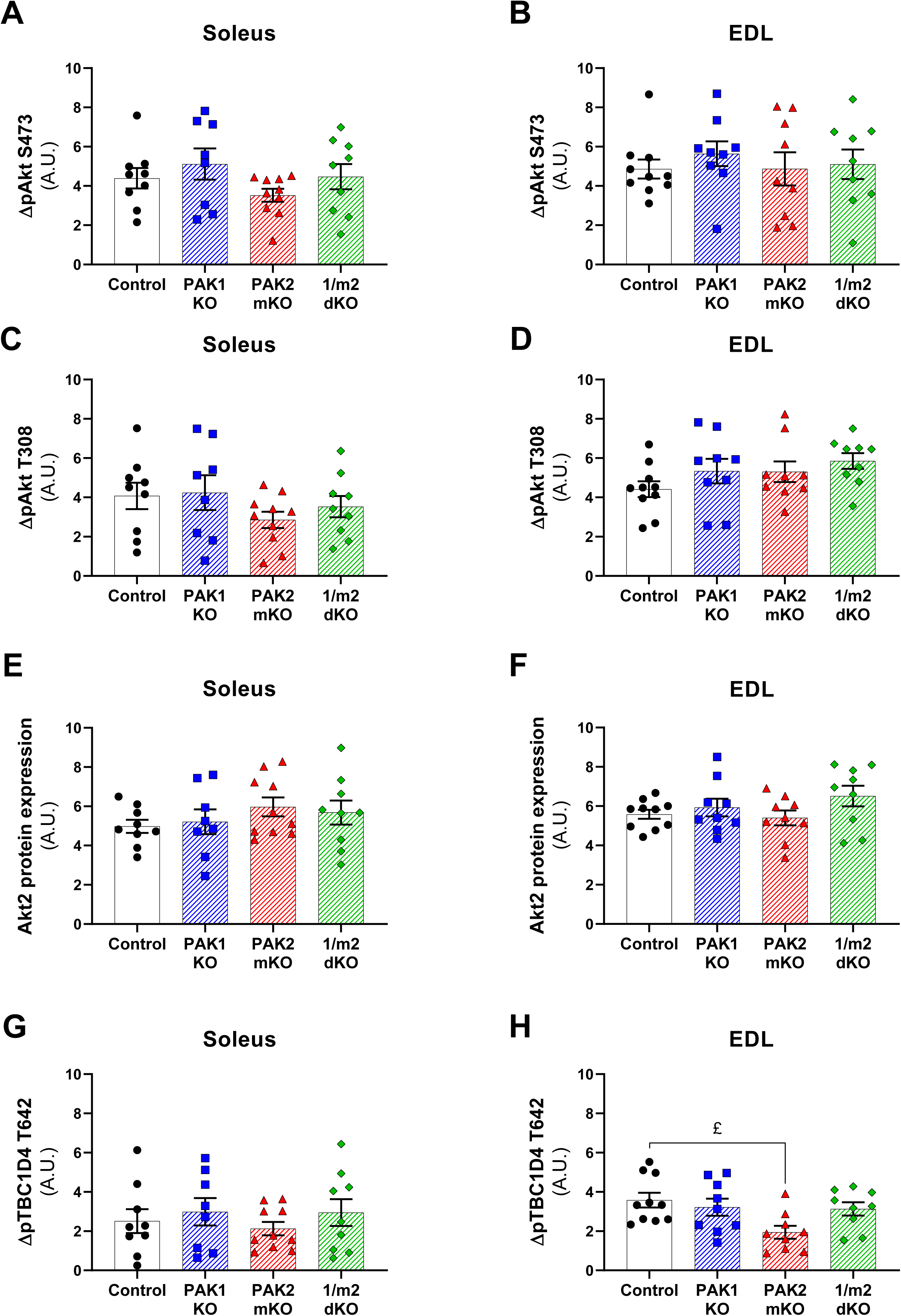
**(A-D)** Δ-phosphorylated (p)-Akt S473 and ΔpAkt T308 in insulin-stimulated (60 nM) soleus (A and C) and extensor digitorum longus (EDL; B and D) muscle from whole-body PAK1 knockout (KO), muscle-specific PAK2 (m)KO, PAK1/2 double KO (1/m2 dKO) mice or control littermates from Fig. 7A-D. **(E-F)** Quantification of total Akt2 protein expression in soleus (E) and EDL (F) muscle from whole-body PAK1 KO, PAK2 mKO, 1/m2 dKO mice or control littermates. Total protein expression is an average of the paired basal and insulin-stimulated sample. **(G-H)** ΔpTBC1D4 T642 in insulin-stimulated (60 nM) soleus (G) and EDL (H) muscle from whole-body PAK1 KO, PAK2 mKO, 1/m2 dKO mice or control littermates from Fig. 7G-H. Statistics were evaluated with a two-way ANOVA to test the factors ‘PAK1’ (PAK1+/− vs. PAK−/−) and ‘PAK2’ (PAK2^fl/fl^;MyoD^+/+^ vs. PAK2^fl/fl^;MyoD^iCre/+^) thereby assessing the relative contribution of PAK1 and PAK2, respectively. Differences between genotypes were evaluated with a one-way ANOVA. The number of determinations in each group: Control, *n = 9/10* (soleus/EDL); PAK1 KO, *n = 8/9*; PAK2 KO, *n = 10/9*; 1/m2 dKO, *n = 9/9*. Significant one-way ANOVA was evaluated by Tukey’s post hoc test: Control vs. PAK2 mKO £ (p<0.05). Data are presented as mean ± S.E.M. with individual data points shown. A.U., arbitrary units.

## Notes

#### Summary of Updates

Addition of new data. Removal of data on the involvement of group I PAKs in contraction-stimulated glucose uptake which will be included in a different manuscript.

